# Proteomic evidence for amyloidogenic cross-seeding in fibrinaloid microclots

**DOI:** 10.1101/2024.07.16.603837

**Authors:** Douglas B Kell, Etheresia Pretorius

**Affiliations:** Department of Biochemistry, Cell and Systems Biology, Institute of Systems, Molecular and Integrative Biology, Faculty of Health and Life Sciences, University of Liverpool, Crown St, Liverpool L69 7ZB; The Novo Nordisk Foundation Centre for Biosustainability, Building 220, Søltofts Plads 200, Technical University of Denmark, 2800 Kongens Lyngby, Denmark; Department of Physiological Sciences, Faculty of Science, Stellenbosch University, Stellenbosch Private Bag X1 Matieland, 7602, South Africa

**Keywords:** clotting, amyloid, fibrinaloid, proteomics, cross-seeding, fibrils

## Abstract

In classical amyloidoses, amyloid fibres form through the nucleation and accretion of protein monomers, with protofibrils and fibrils exhibiting a cross-β motif of parallel or antiparallel β-sheets oriented perpendicular to the fibre direction. These protofibrils and fibrils can intertwine to form mature amyloid fibres. Similar phenomena occur in blood from individuals with circulating inflammatory molecules (also those originating from viruses and bacteria). In the presence of inflammagens, pathological clotting can occur, that results in an anomalous amyloid form termed fibrinaloid microclots. Previous proteomic analyses of these microclots have shown the presence of non-fibrin(ogen) proteins, suggesting a more complex mechanism than simple entrapment. We provide evidence against a simple entrapment model, noting that clot pores are too large and centrifugation would have removed weakly bound proteins. Instead, we explore whether co-aggregation into amyloid fibres may involve axial (multiple proteins within the same fibril), lateral (single-protein fibrils contributing to a fibre), or both types of integration. Our analysis of proteomic data from fibrinaloid microclots in different diseases shows no significant overlap with the normal plasma proteome and no correlation between plasma protein abundance and presence in microclots. Notably, abundant plasma proteins like α-2-macroglobulin, fibronectin, and transthyretin are absent from microclots, while less abundant proteins such as adiponectin, periostin, and von Willebrand Factor are well represented. Using bioinformatic tools including AmyloGram and AnuPP, we found that proteins entrapped in fibrinaloid microclots exhibit high amyloidogenic tendencies, suggesting their integration as cross-β elements into amyloid structures. This integration likely contributes to the microclots’ resistance to proteolysis. Our findings underscore the role of cross-seeding in fibrinaloid microclot formation and highlight the need for further investigation into their structural properties and implications in thrombotic and amyloid diseases. These insights provide a foundation for developing novel diagnostic and therapeutic strategies targeting amyloidogenic cross-seeding in blood clotting disorders.

## Introduction

Blood homeostasis is a finely tuned process involving a complex series of reactions known as the blood clotting cascade. Central to this process is the conversion of fibrinogen to fibrin, catalysed by the serine protease thrombin, resulting in the formation of polymeric fibrin clots. Fibrinogen, a major plasma protein, is pivotal in this cascade, undergoing a remarkable transformation from a soluble protein to an insoluble fibrin matrix. While the mechanisms of normal clot formation are well understood, recent studies have unveiled the formation of pathological clot structures, termed fibrinaloid microclots, in the presence of inflammatory molecules, including those with viral and bacterial origins. These fibrinaloid microclots display unique proteomic profiles and amyloid-like properties, suggesting a complex interplay between protein misfolding and aggregation beyond simple entrapment. This paper explores the proteomic characteristics of fibrinaloid microclots, investigates the mechanisms of their formation, and highlights their potential implications in thrombotic and amyloid diseases.

An important part of blood homeostasis involves the blood clotting cascade. This is well known (Figure 1A) and our focus is on the catalysis by the serine protease thrombin of the conversion of fibrinogen to make polymeric fibrin. Fibrinogen is one of the most abundant plasma proteins, present at some 2-4 g.L^-1^ (e.g. [1–4]). It is a cigar-shaped molecule of ca 5 x 45nm, and consists of several chains (e.g. [5–8]). The action of thrombin leads to the serial removal of two fibrinopeptides, which sets in motion a remarkable self-organisation by which the fibrinogen molecules interact to form protofibrils, **fibrils, and then fibres of some 50-100nm diameter (Figure 1B), implying some hundreds of** fibrinogen molecules in each length element of the typical fibrin fibre. In normal clots the direction of the fibrinogen molecules and fibrin protofibrils is parallel to that of the fibres.

**Figure 1.**
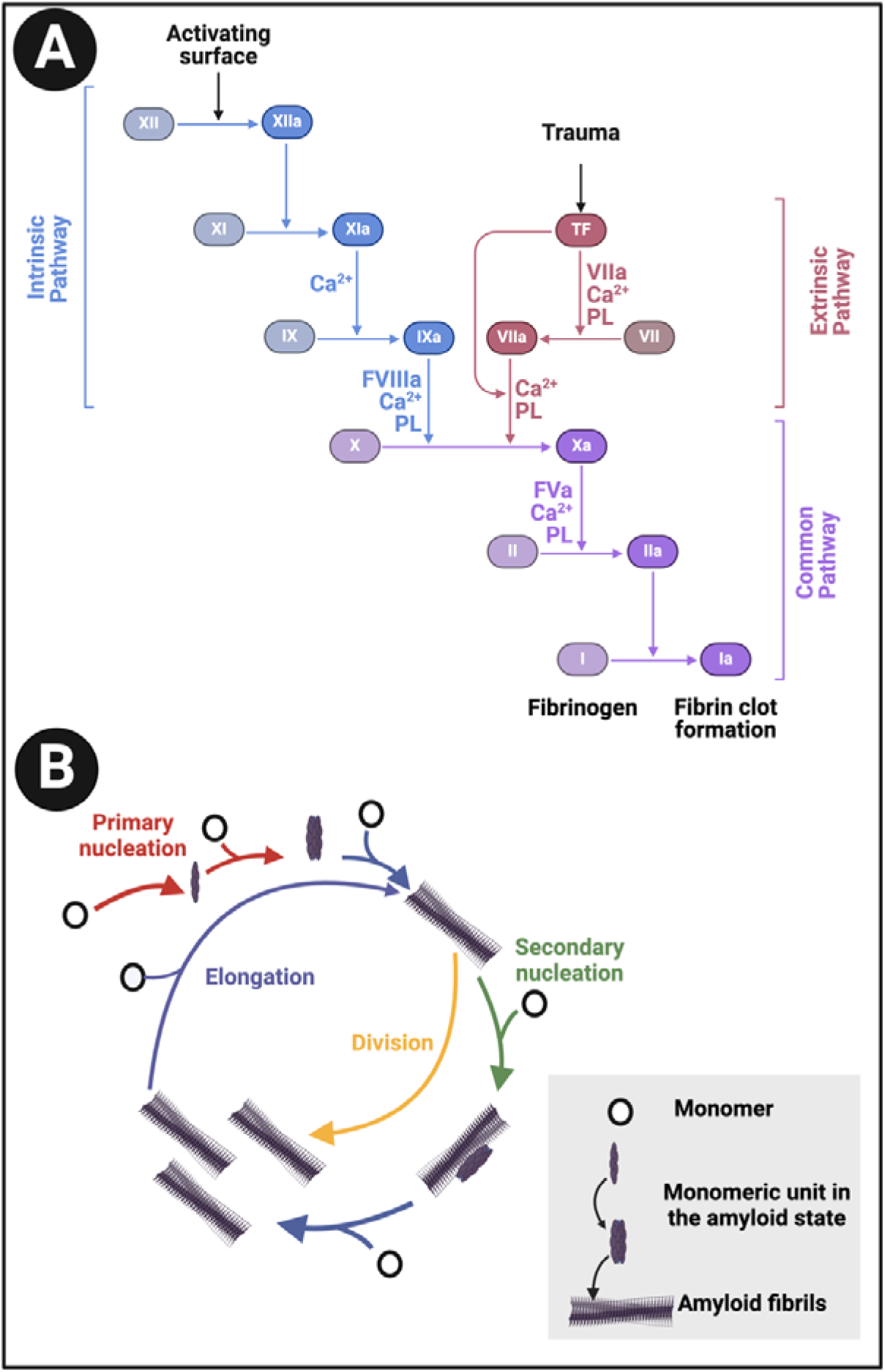
**(A):** The clotting cascade and **(B)** fibrinogen conversion to fibrin. The clotting cascade involves the intrinsic, extrinsic, and common pathways, each comprising various clotting factors. The intrinsic pathway includes factors I (fibrinogen), II (prothrombin), IX (Christmas factor), X (Stuart-Prower factor), XI (plasma thromboplastin), and XII (Hageman factor). The extrinsic pathway consists of factors I, II, VII (stable factor), and X. The common pathway involves factors I, II, V, VIII, and X. These factors circulate in the bloodstream as zymogens and are activated into serine proteases, which catalyze the cleavage of subsequent zymogens into more serine proteases, ultimately activating fibrinogen. The serine proteases include factors II, VII, IX, X, XI, and XII, while factors V, VIII, and XIII are not serine proteases. The intrinsic pathway is activated by exposed endothelial collagen, whereas the extrinsic pathway is triggered by tissue factor released by endothelial cells after external damage. Drawn using Biorender.com.

The pore sizes of typical clots are of the order 0.5-5 μm when fibrinogen is at its physiological concentrations (e.g. [9–13]) (the pore diameters can be far lower at massively extraphysiological fibrinogen concentrations [14]), so without specific binding of some kind they are clearly incapable of simply entrapping molecules of globular proteins (with diameters in the low-nm range). A complex set of reactions also contribute to normal clot degradation (fibrinolysis) [15].

### Proteins of identical sequence can adopt alternative, stable macrostates

The famous ‘unboiling an egg’ experiment of Christian Anfinsen [16, 17] involved the chemical denaturation of ribonuclease followed, upon removal by dialysis of the chemical denaturant, of its refolding into what was considered to be the same native form as the original made naturally following ribosomal synthesis. Importantly, this led to the conclusion that the information necessary for a protein’s tertiary structure could be encoded solely in its primary amino acid sequence. Unfortunately, it was also widely assumed that this structure was thus the thermodynamically most stable under the conditions of interest. This latter assumption could only be just that (an assumption) because of the astronomical number of conformations that a string of n amino acids might adopt even as a single molecule [18, 19], leave alone when ensembles of a given protein form inclusion bodies. Indeed we now know of many classes of example in which proteins can adopt very different conformations despite having the identical primary structure. Prions, amyloids and prionoids are three such classes [20–23]. In fact, the existence of very different macrostates or folds, between which proteins can in fact switch physiologically, is surprisingly common [24–27]. Such proteins have been referred to as ‘metamorphic’ [28–32]. While the present versions of Alphafold cannot (and were not designed to) predict such a multiplicity of macroscopic conformations effectively [33, 34] (though see their utility for isoenergetic microstates [35]), the primary sequences do in fact contain such information [36, 37]. There is also evidence that proteins capable of adopting multiple macrostates were in fact selected adaptively [38–41], and can be designed accordingly [42].

Given the above, and that our focus here is on amyloidogenic proteins, it is of special interest that α-helix-to-β-sheet transitions are a noteworthy feature of such proteins [36], and particularly, for our present purposes, some in SARS-CoV-2 [43], where several proteins are amyloidogenic [44, 45]. We note too that some alleles of the fibrinogen Aα chain may produce highly amyloidogenic proteolytic fragments [46], and that fibrinogen can bind to well-established amyloids such as Aβ [47–50].

### Structure and interconversion of amyloid(ogenic) proteins

In contrast to the expectations of the Anfinsen experiment, it has become well established that many (and maybe even most) proteins can fold into states that are considerably more stable than the one natively or most commonly adopted as they leave the ribsome, but that this thermodynamically favoured conformation (or, more accurately, the set of isoenergetic conformations) is normally kinetically inaccessible due to a massive energy barrier of some 36-38 kcal.mol^-1^ [21, 51]. A particular class of these more stable conformations involve a cross(ed)-β-sheet motif [52–58], and they become insoluble because they tend to aggregate and self-assemble; following their discovery by Virchow in 1854 (reviewed by Sipe and Cohen [59]) they are referred to as amyloids (see e.g. [60–65]). As is well known, they are intimately (if at best only partially) involved in a variety of diseases, including Alzheimer’s [66] and Parkinson’s. Such syndromes are collectively referred to as amyloidoses (e.g. [67–73]. As phrased by Burdukiewicz *et al*. [74], “Despite their diversity, all amyloid proteins can undergo aggregation initiated by short segments called hot spots.” This is a massive field, so our focus is on those parts that most reflect the present core question, which is around their self-assembly. This – commonly the transition from an α-helix structure to a β-sheet one [75] – necessarily involves partial unfolding [76, 77].

### Rules for amyloidogenesis and cross-β formation

In contrast to the classical secondary structure of β-sheets, where the rules for their formation in terms of amino acid sequence are broadly polar-apolar-polar-apolar-(etc) [78], the sequence rules for amyloidogenic potential generally, and for cross-β sheet formation [79] in particular, are rather more obscure. This is not helped by the relative paucity of data on (sub)sequences that can encode amyloidogenicity, but experimental and computational progress is being made (e.g. [80–82] and Table 1) and databases of amyloidogenic hexapeptides [83] and amyloid-amyloid interactions exist [84]. It is indeed reasonable that the most predictive properties for residue amyloidogenicity are “hydrophobicity index [85], average flexibility indices (a normalized fluctuational displacement of an amino acid residue) [86], polarizability parameter [87] and thermodynamic β-sheet propensity [88]” [74].

**Table 1.**
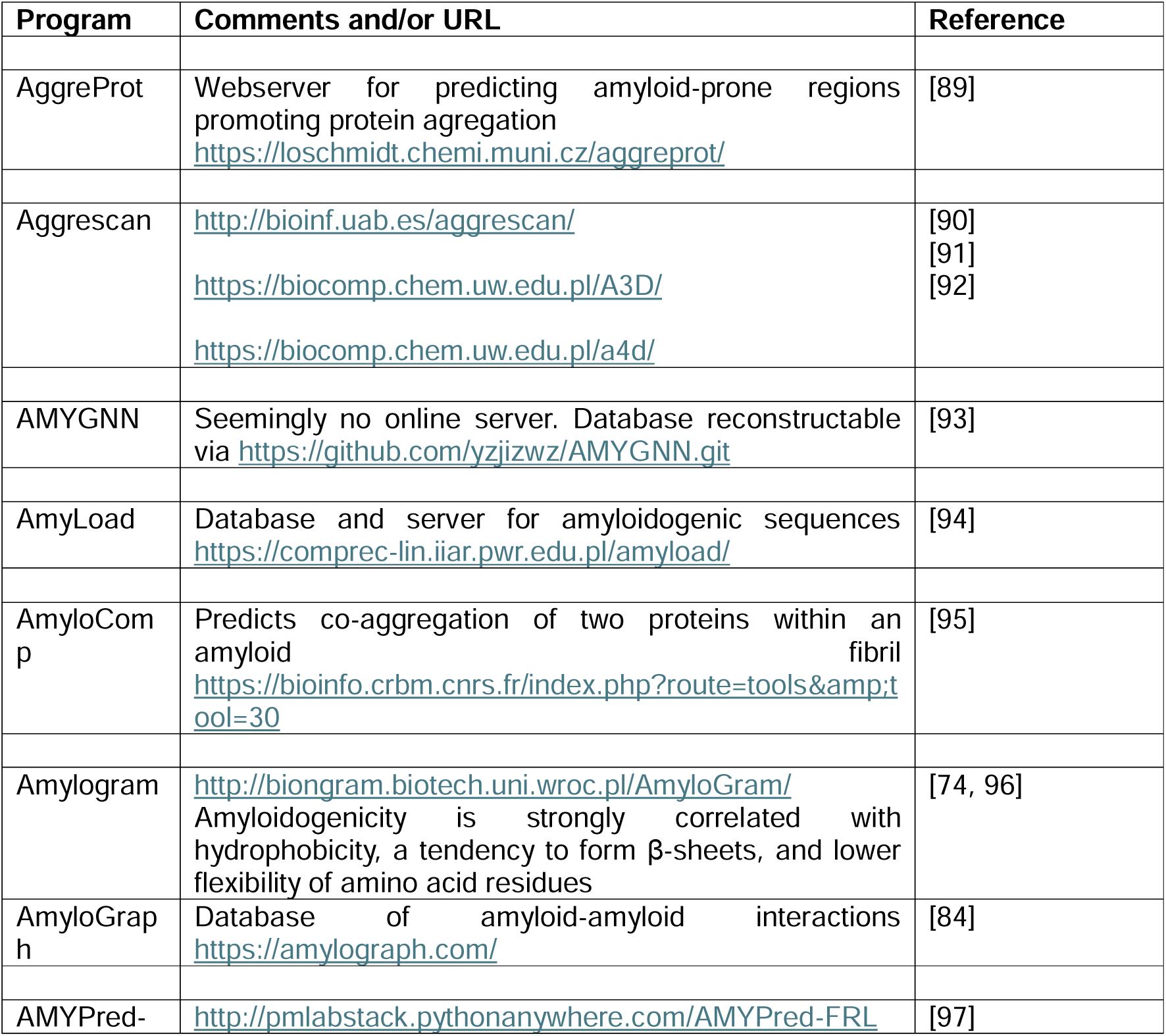
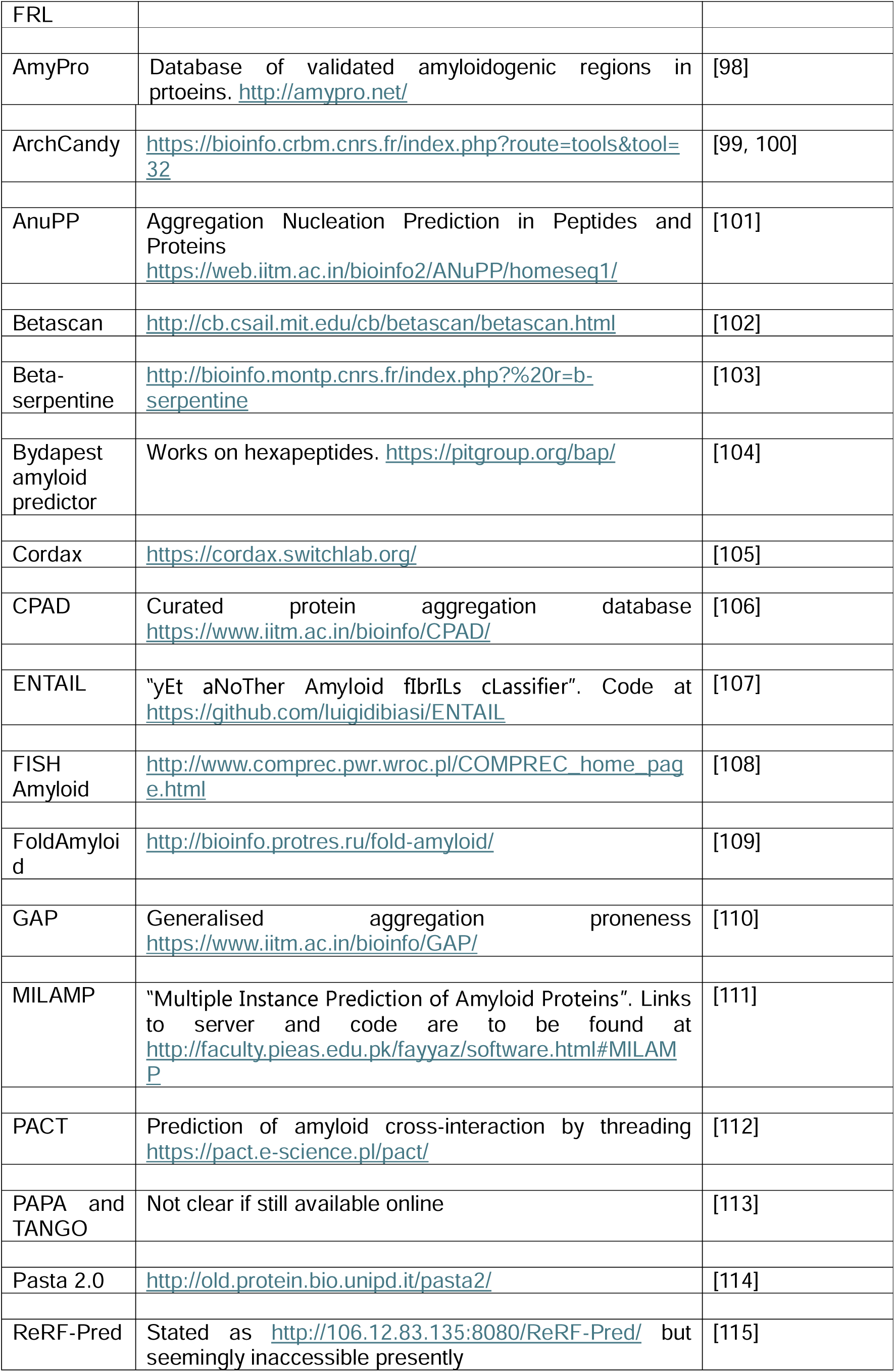
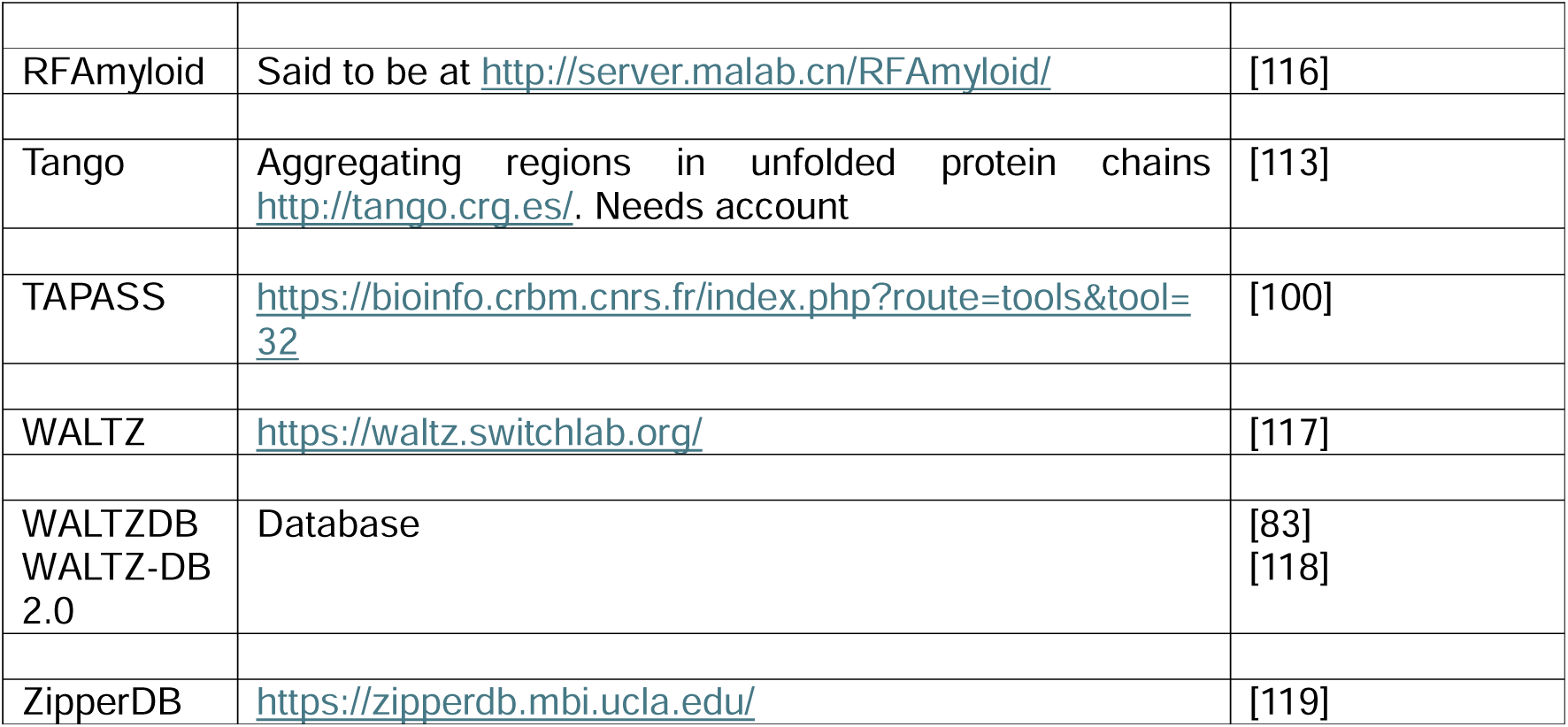
Some computational resources for predicting amyloidogenic regions in proteins.

Figure 2A shows a prediction from AmyloGram [74] of a well-established amyloidogenic protein in the form of the human prion protein PrP^c^, when a significant run of residues towards the C-terminus has an amyloidogenicity score (referred to as a ‘probability of self-assembly’) exceeding 0.75 (and see later). Figure 2B also shows the predictions for the fibrinogen A/α chain, which shares some of these features. Figure 2C shows the predictions for the fibrinogen A/α chain at AnuPP, giving a broadly similar picture.

**Figure 2.**
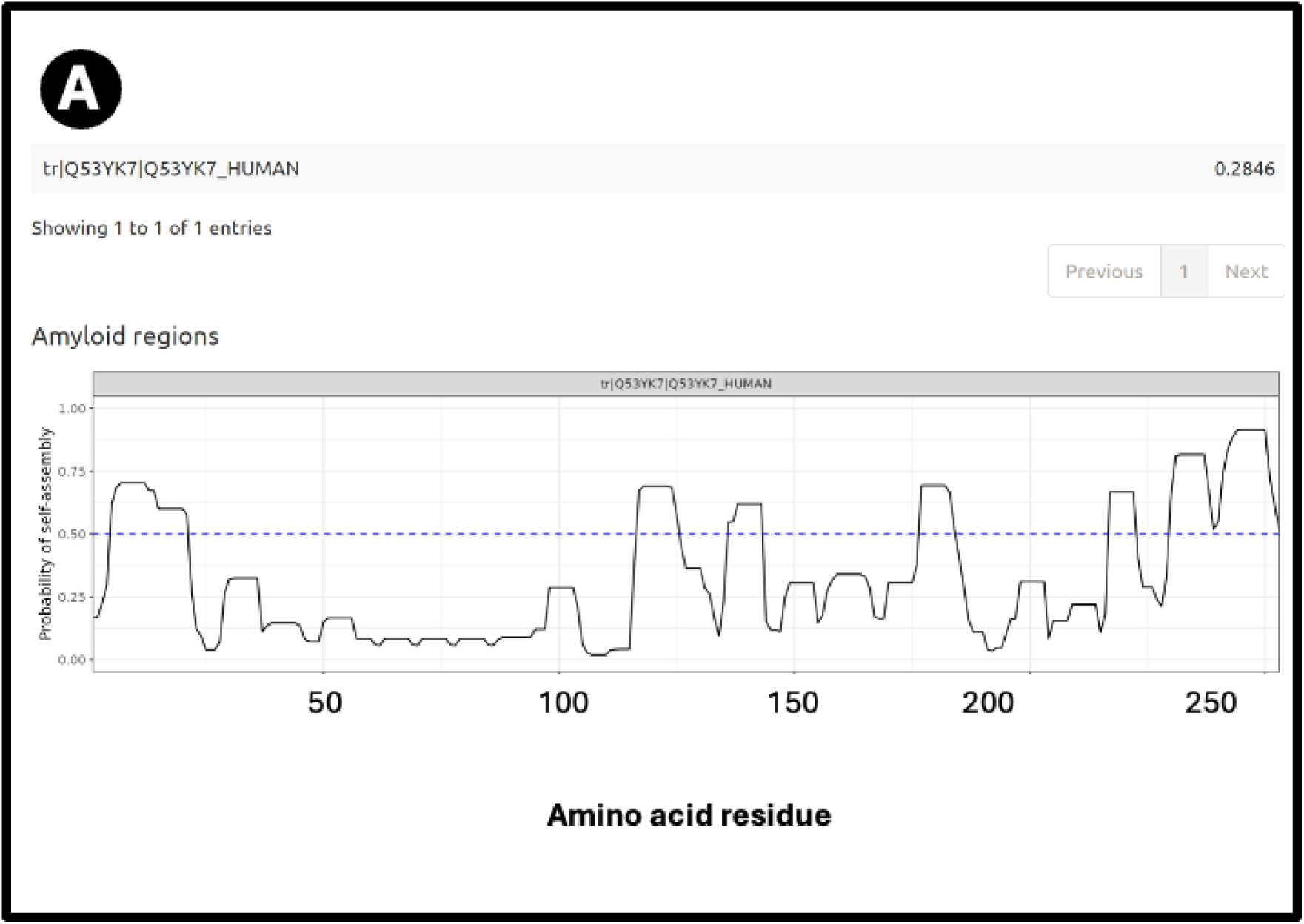
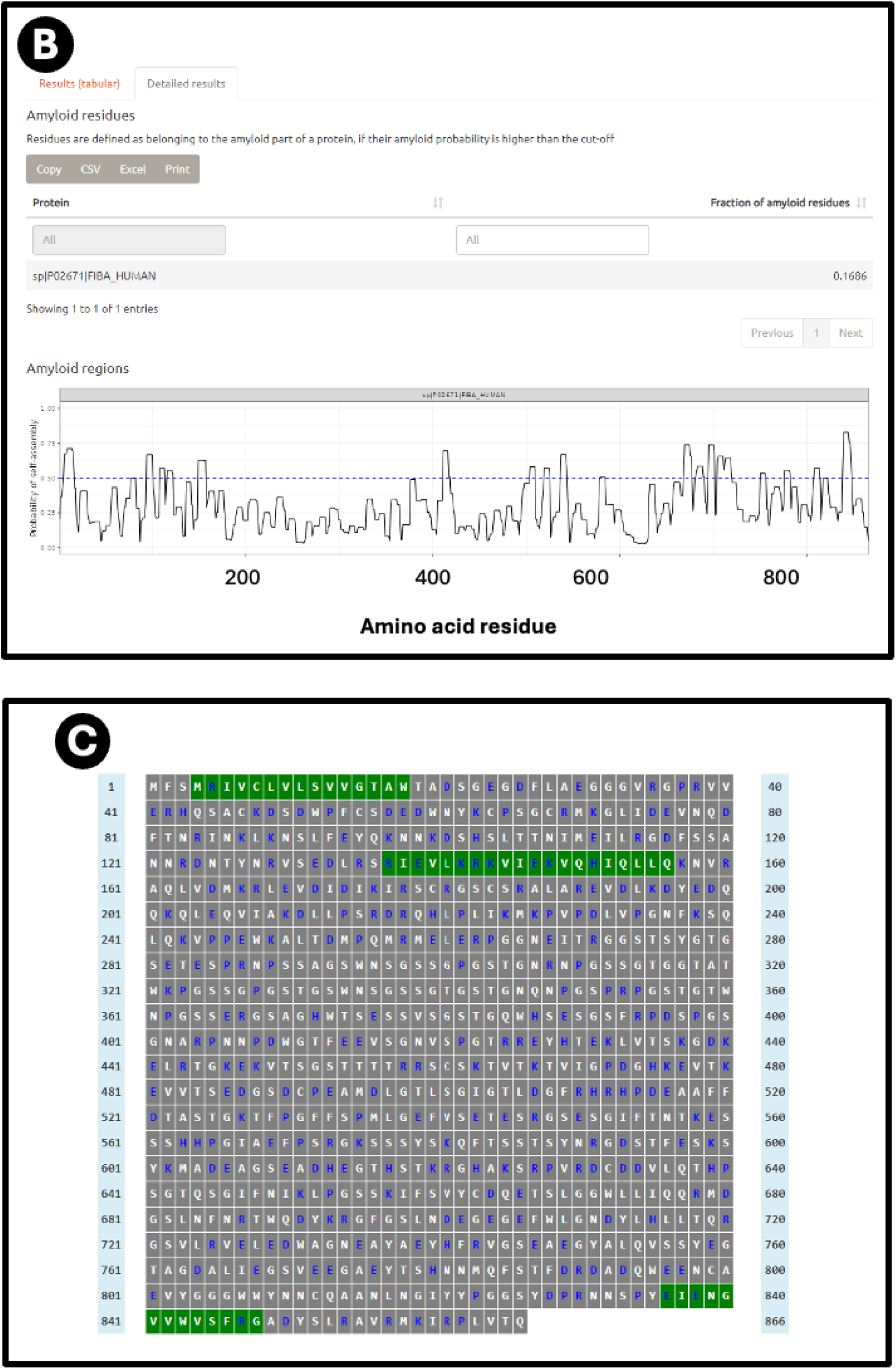
Prediction of amyloidogenic regions of **(A)** human prion protein and **(B)** fibrinogen α chain at AmyloGram [74, 96], and **(C)** fibrinogen α chain at AnuPP [101]. In the latter case amylodogenic regions are shown in green.

### Prevalence of amyloidogenicity

As can be established by testing various sequences on the above servers (we focused on AmyloGram [74, 96], see later), as well as the classical amyloidoses, a very great many [120–122] (and possibly most [123]) proteins can exhibit amyloid formation under certain conditions [65, 124]. Examples include insulin [125–127], lysozyme [128–136], proteins providing structure/texture in various processed foods and other gels [137–139], and even certain yeast [140–142] and bacterial [143] proteins, including some that can be ‘inherited’.

### Amyloid structures

Historically, establishing the structures of amyloid fibres formed even by single proteins or peptides was difficult because of their insoluble nature, but this is being changed by techniques such as solid-state NMR (e.g. [144–148]) and nowadays, in particular, cryoEM (e.g. [149–154]). These make it clear that amyloidogenic stretches of proteins can be responsible for cross-β fibril and fibre formation. An example is given in Figure 3, reproduced from an Open Access paper [154].

**Figure 3:**
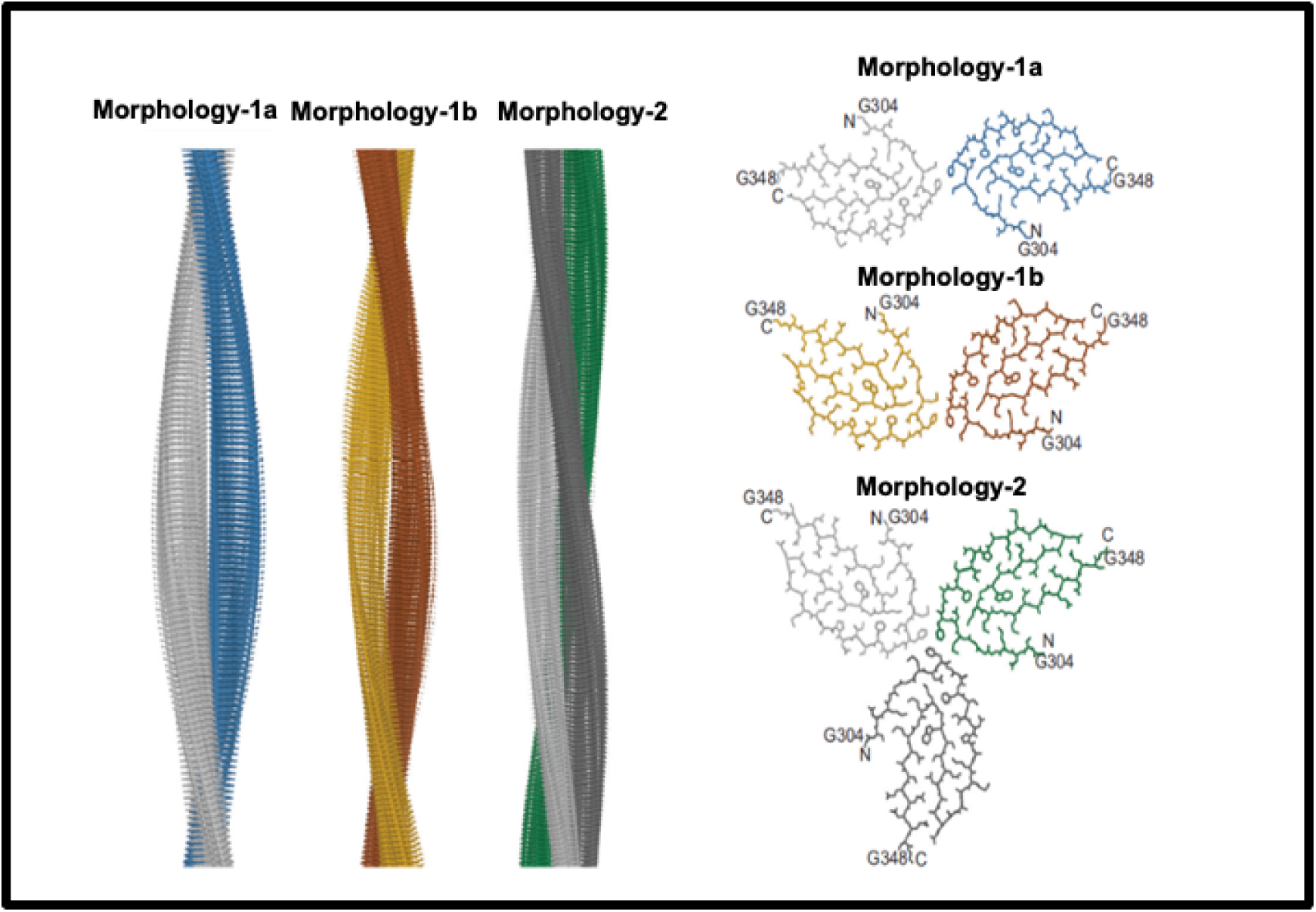
Structures of Amyloid fibres: Fibril formation by cross-ß elements. Reproduced from an Open Access paper [154].

### Amyloid detection with thioflavin T and other stains

A continuing and historically important discovery was the fact that the dye thioflavin T (ThT; https://pubchem.ncbi.nlm.nih.gov/compound/Thioflavin-T) binds to a whole series of amyloids, with a concomitant increase in its fluorescence [155]. This occurs because rotation of the normally rotatable single bond between the benzothiazole and dimethylaniline rings allows fluorescence from an excited state to be dissipated and hence quenched. When the ThT is bound appropriately to a macromolecule, no such rotation is possible, fluorescence occurs, and thus ThT is a fluoro<u>genic</u> stain for amyloids (e.g. [156–163]). Note too that in the absence of amyloid target, ThT forms micelles with a critical micelle concentration of some 4 μM [156]. As phrased by Biancalana and Koide [158], “ThT binds to diverse fibrils, despite their distinct amino acid sequences, strongly suggesting that ThT recognizes a structural feature common among fibrils. Because amyloid fibrils share the cross-β architecture, it is generally accepted that the surfaces of cross-β structures form the ThT-binding sites”. Indeed a number of such crystal structures have been solved, e.g. [164], or the relevant structures determined by other means (e.g. [165–168]), binding being seen as perpendicular to the cross-β elements and thus parallel to the fibrils themselves [52, 144, 166]. Figure 4 gives an example from an Open Access publication [169].

**Figure 4:**
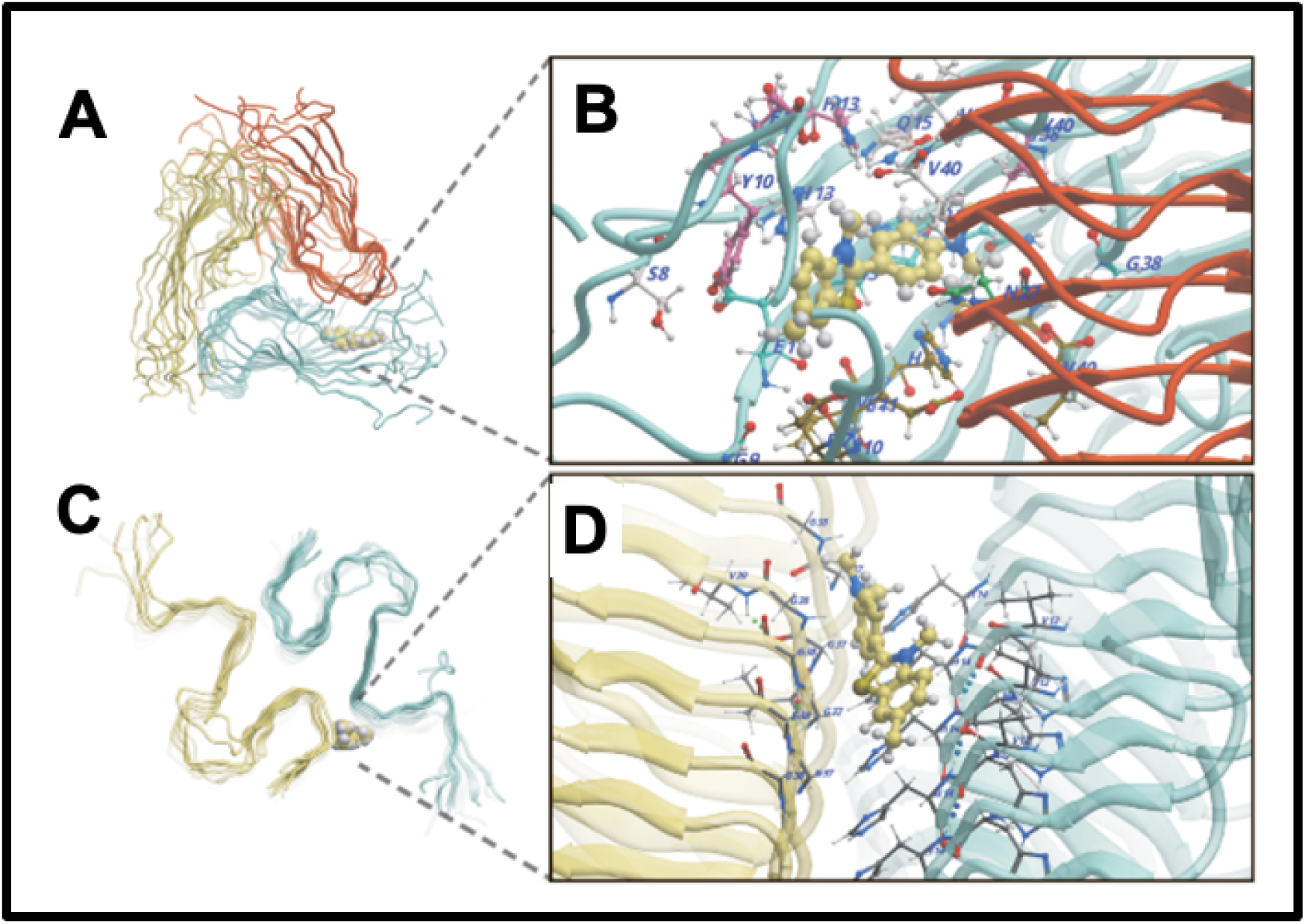
Thioflavin T (ThT) binds to amyloid fibrils by recognizing a structural feature common among them. Amyloid fibrils share a cross-β architecture, where the β-sheets are oriented perpendicular to the fibril axis. The surfaces of these cross-β structures form the binding sites for ThT, which results in a characteristic increase in fluorescence upon binding. This property makes ThT a widely used fluorescent stain for detecting and studying amyloid fibrils.

Note, however, that there can be quite subtle differences in the binding modes of ThT to specific amyloids, leading to changes in fluorescence intensity [169–171]. All of this said, and noting that ThT can in some cases bind to non-amyloid structures (e.g. [172]), ThT certainly remains the most popular amyloidogenic dye (e.g. [21, 157, 158, 161, 162, 173, 174]).

This difference in the precise mode of binding of a given dye can be observed in the studies of oligothiophene dyes (marketed as ‘Amytrackers’) by Nilsson and colleagues, where spectral as well as intensity changes can be observed between different amyloid structures (e.g. [175–180]).

Many other fluorogenic dyes that bind amyloid also exist, some with desirable spectral properties. Examples include NIAD-4 [181] that has an enhanced Stokes shift [163, 182–186], and others that excite and emit towards the (near infra)red end of the spectrum (e.g. [184, 187–190]).

### Alternative blood clotting

In the same way that many proteins can adopt an amyloid form, as described above, it was discovered [191, 192] that fibrinogen can polymerise into an anomalous amyloid-like form, that stained very strongly with thioflavin T and indeed other stains such as Amytracker stains (see Figure 5 of confocal micrographs where ThT and Amytrackers were exposed to plasma from healthy participants and those with type 2 Diabetes [193]. The typical size of these clots, that were termed microclots (e.g. [194–198]) is in the range 2-200 μm (e.g. [191, 195, 198–202])

**Figure 5:**
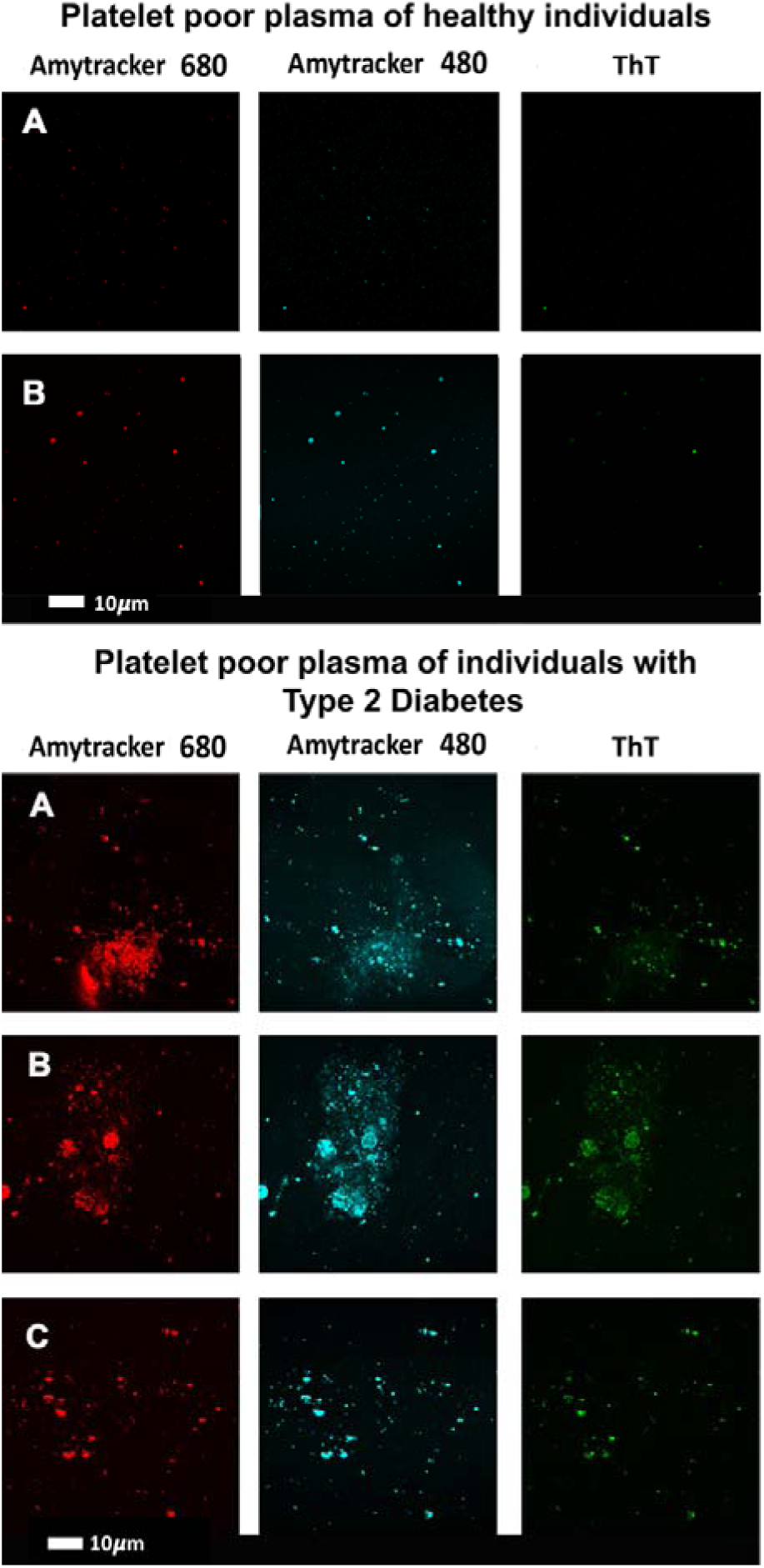
Confocal micrographs where thioflavin T (T) and Amytrackers were used to stain plasma from healthy participants and those with type 2 Diabetes [193].

Acording to AmyPro [98] the most amyloidogenic region of fibrinogen α chain encompasses residues 148-160 (KRLEVDIDIKIRS), though Amylogram implies a second region nearer the C-terminus is even greater (Figure 2A).

It is interesting to note that fibrin<u>ogen</u> is itself able to interact with other small amyloids, actually inhibiting their extension into larger fibrils [203–205], indicating that while it is well capable of binding amyloidogenic sequences, its own amyloidogenicity is only normally manifest <u>during clotting itself</u>, under the action of thrombin, though certain amyloidogenic alleles can lead to a fibrin amyloidosis (e.g. [206–208]). Other peptides can bind preferentially to (at least normal forms of) fibrin but not fibrinogen [209–211].

### Size of fibres in classical amyloidoses and in normal and fibrinaloid clotting

To assist in understanding the nature of the fibres and how they may differ between ‘normal and fibrinaloid microclots, it is worth rehearsing the diameter of a typical monomer fibril of a cross-β element in a fibril. This depends, of course, on the length of the amyloidogenic run of amino acids that forms it, but is typically 1-2nm or so. A protofibril consisting of 2-4 intertwined monomer fibrils may be 4-11nm for molecules such as Aβ [212], 7nm for tau [213], 11nm for the prion protein in its amyloid PrP^Sc^ form [214], 6-15 nm for α-synuclein [215, 216], and 7-13 nm for transthyretin [217].

By contrast (though cf. [96] for artificial super-amyloidogenic hexapeptides), the diameter of individual clot fibres is roughly 100 nm for amyloid clots (e.g. [191]) and is similar in many cases for normal ones [218], but can be as much as 400 nm or even more for normal, non-amyloid ones [219–223]. For normal non-amyloid clots this would require several hundred elements, and for the amyloid version very long runs of crossed-β features (that run in a criss-cross manner perpendicular to the long axids of the fibre, and many, many protofibrils intertwining around each other by lateral co-aggregation.

Normal clots are far better studied, and their diameter, for instance, depends on the fibrinogen concentration, consistent with general chemical kinetics. However, what the exact structures are, especially for the fibrinaloid ones, and what eventually stops them increasing in both length and diameter indefinitely, is not yet known. The ability of normal fibrinogen [224] and other proteins [225] to convert into a β-sheet-rich form is probably highly relevant. These observations also depend, of course, on a variety of factors such as degree of hydration, initial fibrinogen [226] and thrombin [227] concentrations, levels of small molecules [191, 228], of metal and other ions [229–232], and so on. However, the point of this paragraph is that these are clearly very much larger numbers for the fibre diameters in fibrinaloid microclots than are those seen in classical amyloidoses.

### Inclusion bodies, compared with the growth and aggregation of classical amyloid fibrils

Inclusion body formation is a well-known feature of recombinant protein expression (e.g. [233–240]), and is usually considered to occur due to the protein it is desired to fold being unable to keep up with the rates of its synthesis. Inclusion body formation largely involves a somewhat random or amorphous type of aggregation driven by interactions between hydrophobic residues of proteins that have failed to fold properly, even if they may sometimes contain or induce amyloid-like structures [124, 241–247]. They mostly consist of the same polypeptide (so are sometimes considered a useful means of recombinant protein purification) but can certainly entrap other proteins via non-covalent interactions [245, 248]

This contrasts with the type of ordered self-organisation seen in amyloid fibrils where multiple copies of the same protein also come together but into much more regular or structured shapes (e.g. [61, 76, 77, 249–254]). The hallmark is one of various parallel or antiparallel cross-β sheet motifs [46, 255–258] that run perpendicular to the fibril axis. They provide for a very characteristic X-ray diffraction peak reflecting a spacing of some 4.7Å [259].

A given protein can even adopt various amyloid forms, known here as polymorphisms (e.g. [52, 260–271]). The same is true of prions (e.g. [272–276]), arguably the most ‘extreme’ forms of amyloid(ogenic) proteins, and indeed the coinage of the term ‘prionoid’ (e.g. [277–282]) reflects this kind of overlap or continuum. In one sense [21]. It is obvious that there must be parallels between the kind of fibril formation that are seen in classical amyloidogenesis (commonly in the range 2-25nm diameter [252, 283–286]) and that seen in both normal and pathological blood clotting (although those fibrils are commonly at least 10x larger in diameter [195], see above), since in both cases fibrils are an observable result. Lengths of fibrils in classical amyloidoses can be 1 μm or so [287]. Consequently, in this section we rehearse what is known of amyloid fibril formation in the classical amyloidoses [252].

### General phases of amyloid fibril formation

Extensive kinetic and imaging studies *in vitro* (e.g. [65, 252, 288–296]), often using ThT, have recognised several stages of amyloid fibre formation, starting with a lag [163, 297] or nucleation phase creating oligomers (the most cytotoxic forms [298, 299]), then an elongation phase in which protofibrils then fibrils are formed, the latter via protofibrils twisting round each other, and ending with a stationary phase in which fibril and plaque formation is complete (or, more accurately, any elongation and fragmentation or inhibition are occuring at the same rates [252, 291, 300, 301]). Such sigmoidal curves are very similar to those of the batch growth of microbes [302]. It is the fibrils in which the cross-β structures are manifest, implying a structural transition whose detailed mechanism is far from clear.

### Protein entrapment in microclots; cross-seeding

Our own work on the proteomics of fibrinaloid microclots has referred to ‘entrapment’ of non-fibrin proteins in the microclots. However, this <u>cannot</u> be a simple entrapment like a fish in a in a mesh net; the pores are far too big and in any case the centrifugation would have washed soluble proteins away from any weak binding or entrapment. Consequently, the ‘entrapment’ must actually be a forcing of the other proteins to become <u>insoluble</u>, likely by making cross-β sheets and thus joining the tightly-bound-but-noncovalent party and be incorporated into the growing amyloid fibrils. In the terminology of Bondarev and colleagues (Figure 6) and see below), these could be either or both of axial and lateral coaggregation. Evidence for this includes the fact that there is no relationship between what is ‘entrapped’ in the fibrinaloid microclots and the normal plasma abundance of proteins (e.g. albumins and transferrin are pretty well the most abundant and mostly do not appear). In some cases anti-proteolytic sybstances such as antiplasmin [303] and α1 antitrypsin (SERPINA1) [202] may also be present in some abundance.

**Figure 6:**
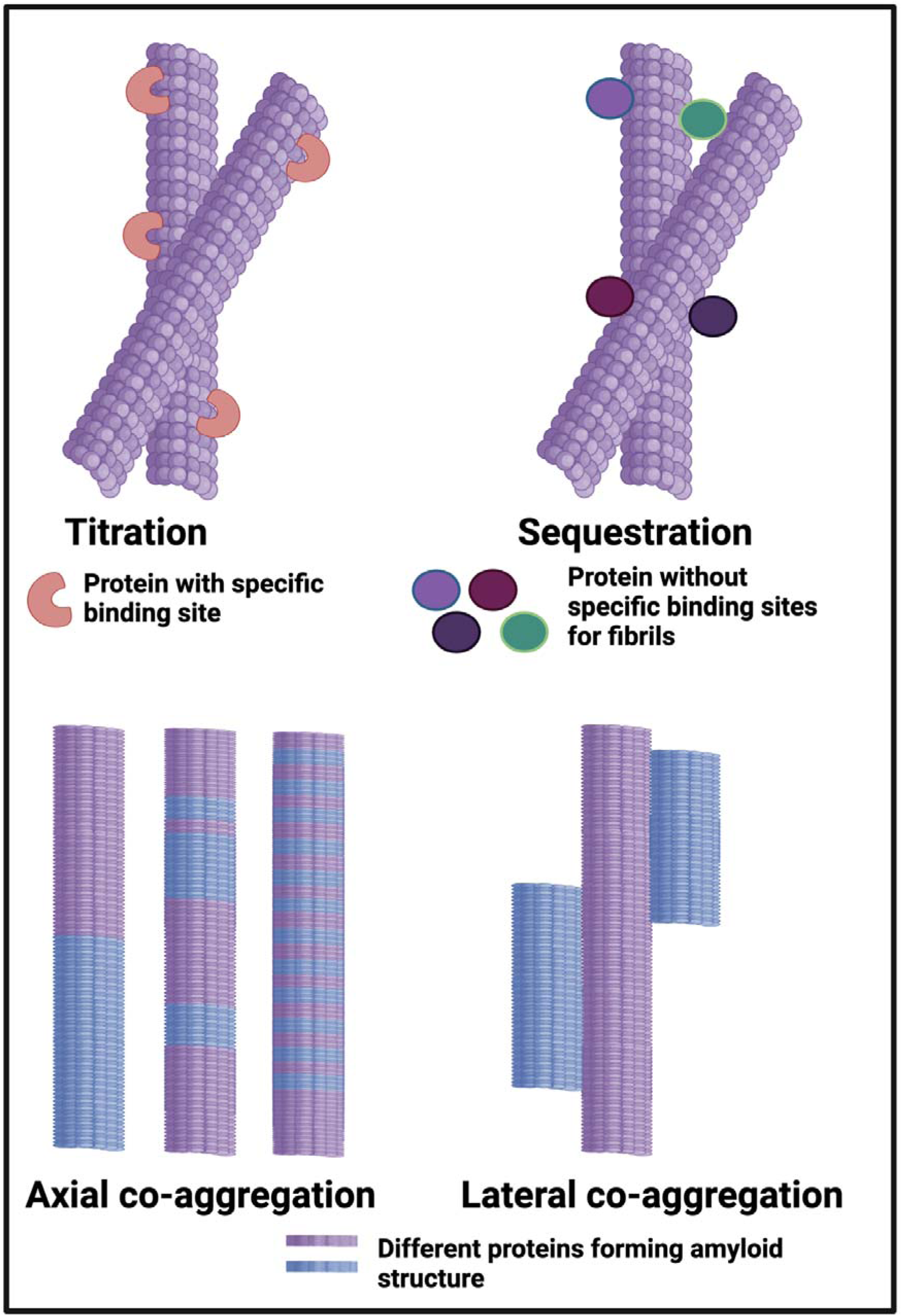
Different classes or types of protein co-aggregation: **(A)** Titration; (**B)** Sequestration; **(C)** Axial and **(D)** Lateral. Adapted from [356].

In a similar vein, many proteins besides α-synuclein are found in the Lewy bodies that can occur in dementia [304, 305], while there is considerable experimental evidence for the co-incorporation of different amyloidogenic proteins into the same fibrils [84, 306–308]. The same is true in transthyretin amyloidosis [309]. This is sometimes referred to as ‘cross-talk’ or ‘cross-feeding’, heterotypic interaction [310, 311], and – perhaps most commonly – ‘cross-seeding’ [169, 312–349]

Note that even simple polyQ features (occuring as C-terminal ‘tails’ in various diseases) can do this [324, 350–354], possibly also by incorporating transition metals [355]. Equually importantly, not all amyloids (‘donors’) can cross-seed other ones [139] (‘acceptors’) and hence be entrapped in the fibrils of the acceptors; there is significant selectivity, whose sequence/structural basis remains unknown.

The occurrence of multiple amyloid proteins within the same fibril is reviewed by Bondarev and colleagues [95, 356], who refer to it as axial co-aggregation (Figure 6). The server AmyloComp [95] (Table 1) also predicts the likelihood of proteins forming axial co-aggregates; that for SERPINA1 and the fibrinogen alpha chain is especially high (unpublished). As phrased by them [95] ‘The core of these amyloid fibrils is a columnar structure produced by axial stacking of β-strand-loop-β-strand motifs called “β-arches” [357–361]’. Well-established examples include RIP1/RIP3 that can induce necroptosis [362] and the HET-s protein that also contains the Rip homotopic interaction motif (RHIM) [363]. Clearly any protein capable of forming these β–arches can then do so so as to make a hetero-fibril, which is what we suggest is the main means of ‘entrapment’ of other proteins in fibrinaloid microclots (provided the amyloidogenic regions are of sufficient length [364]). That fibrinogen <u>can</u> interact with a variety of known amyloidogenic proteins is beyond dispute [365, 366]; causiing them thereby to create new epitopes can even account for autoantibody generation [22].

## Results

### Absence of relationship between microclot proteome and plasma concentration

At least two lines of evidence indicate the lack of relationship between the amount of a protein in plasma and its appearance in fibrinaloid microclots. First, the only overlap between the proteomic data of Kruger and colleagues [367] (who did not report on fibrinogen) and those of Schofield *et al*. [202] was the protein Apolipoprotein A2 (marked in blue in Fig 7). The data were taken from two quite different diseases (acute sepsis [202] vs Long COVID [367]), with ‘normal’ proteome levels spanning several orders of magnitude, and so while the content of these proteins in the average proteomes won’t have differed by more than a factor of two at most, their appearance in the microclots differed massively. Secondly, we extracted the ‘top 20’ data from the pie chart representing the average of three individuals in Figure 3 of [202] and related those (where available) to the average plasma protein concentration [368], indicating that there was no such relation (r^2^ = 0.1 for the data in Fig 7). We also assessed some of the most abundant plasma proteins as tabulated in [369] for their presence or otherwise in the microclots in either study [202, 367] (Figure 8); it is obvious that many of the most abundant proteins are not notably entrapped in the microclots, so those that are are clearly selected, presumably by integration into the amyloid mixtures. (The data underpinning all these analyses are given in a spreadsheet in Supplementary Information.)

**Figure 7:**
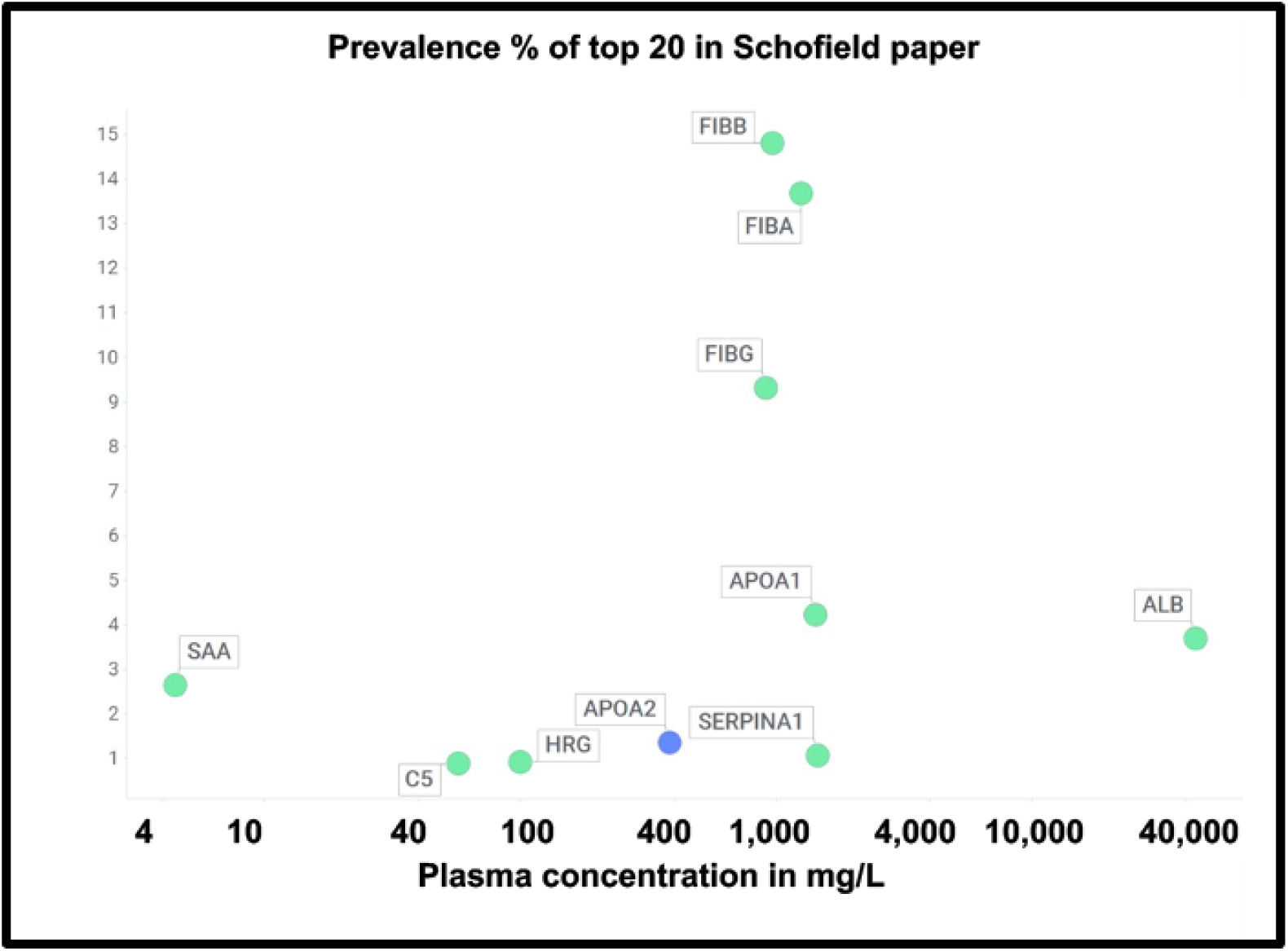
Prevalence of proteins in fibrinaloid microclots in the Schofield ‘top 20’ (green) and the one example also seen in the Kruger study (blue) versus average plasma concentrations that are taken from [368] except for TGFB1 [370] and periostin [371]. Abbreviations as in the List of abbreviations. The line of ‘best fit’ is not shown as it has a correlation coefficient r^2^ of only 0.1.

**Figure 8:**
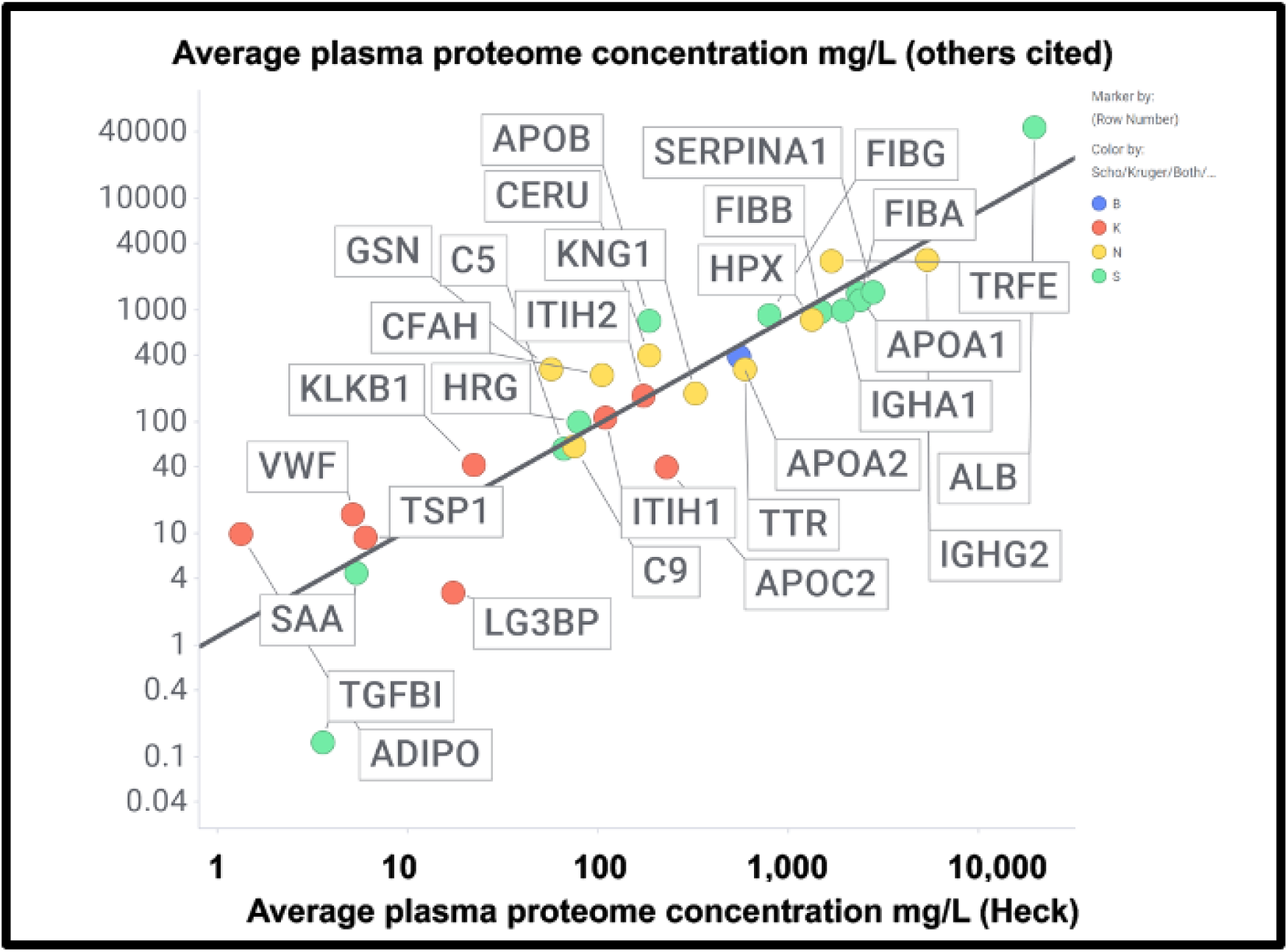
Good correlation between the plasma proteome concentration data in [381] by Heck and colleagues compared with other measurements of the proteome cited in the text and in Supplementary Information, The slope of the line is 0.95 and the correlation coefficient 0.83. Colours encode the datasets in which fibrinaloid proteins were or were not observed, as in Fig 7 and 9, viz Blue both, green Schofield, Red Kruger, Yellow neither. Abbreviations as in the list of abbreviations.

A chief source of protein abundances in plasma is [368]. In addition we used other sources for high-abundance and other detected proteins including C9 [372, 373], Complement Factor H [374], thyroxine-binding globulin [375], retinol-binding protein [376], TGFβ1 [370], periostin [371], CXCL7 (PFA4) [377], (pre)kallikrein [378], galectin-3-binding protein (LG3BP) [379], thrombospondin-1 [380], ITIH1/2 [381]. LG3BP is of interest, as it is substantially lowered in the plasma of those with mild cognitive impairment or Alzheimer’s [382], arguably because it has been removed in amyloid microclots. Similarly, thrombospondin-1 also interacts with Aβ [383]. Reference [381] is very valuable in its own right, since while its coverage lacks some of the low-abundance proteins of interest here, it does list quantative values for 197 plasma proteins; where both are available the concentrations are well correlated (Fig 9)(slope = 1.06, r^2^ = 0.87), taking as the values for the Heck study [381] the averages of six data points for the controls of two patients at the first three time points. Consequently we use the Heck dataset in most of what follows.

**Figure 9:**
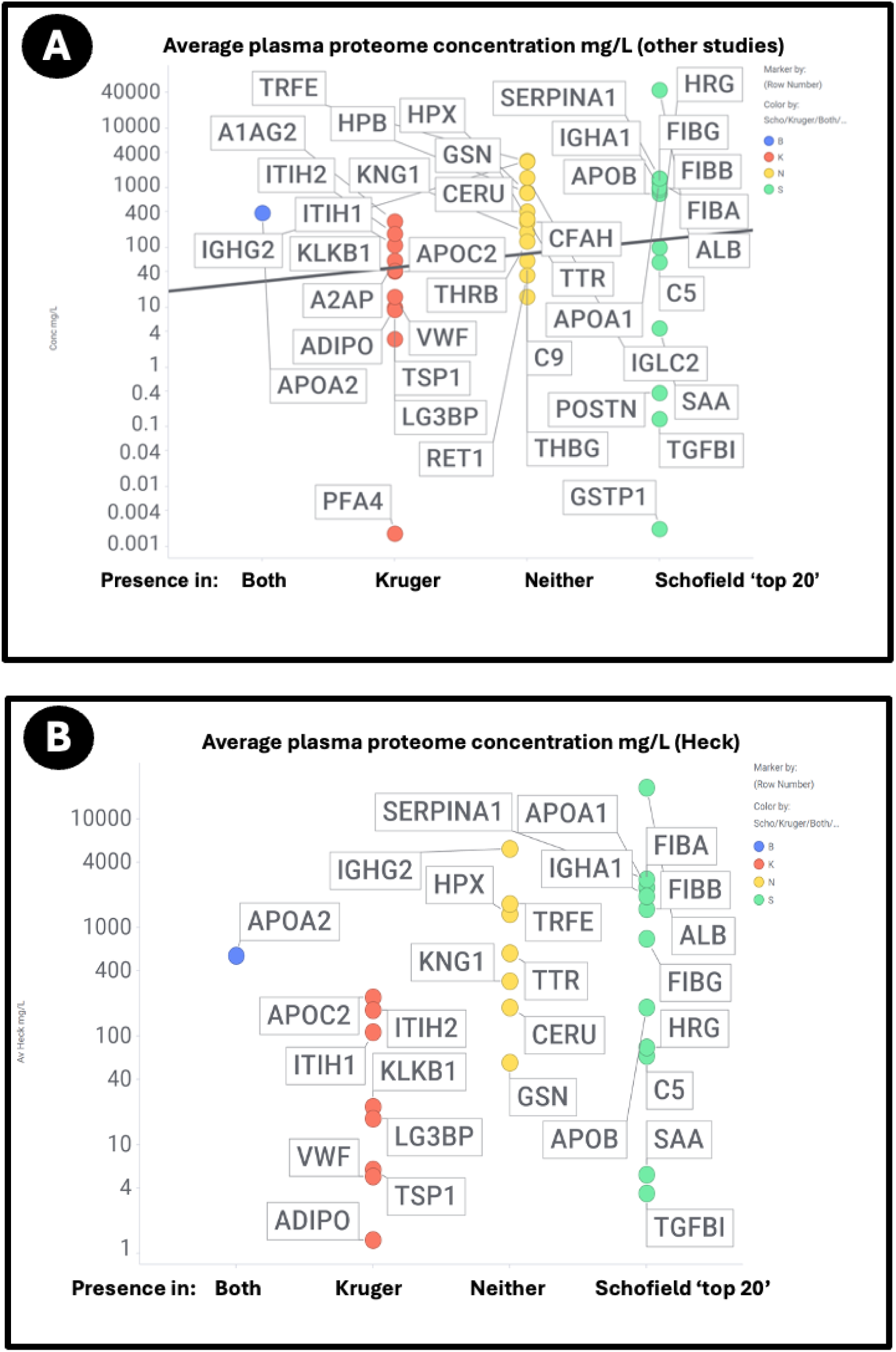
Relationship between the standard plasma proteome concentrations (taken from [381]) and their detection in the Kruger (K), Schofield (S), Both (B) studies or neither (N). Size of symbol encodes protein length in residues. Abbreviations as in list of abbreviations. (**A**) protein concentrations from other studies delineated in the text and the supplementary spreadsheet. (**B**) Proteins from the study of Heck and colleagues [381].

Using both sets of plasma proteome data (since some important markers do not appear in both), we again see large number of proteins that are high in abundance in the plasma proteome that nevertheless are not ‘entrapped’ in the fibrinaloid microclots, and similarly others in low abundance in the proteome nonetheless appear in the fibrinaloid microclots. The conclusion is very clear: there is a significant selectivity with regard to proteins that are entrapped within fibrinaloid microclots. We note too that atomic force microscopy [384] has provided evidence for both axial [385] and lateral [386] co-aggregation, and that cross-seeding can also explain the comorbidities seen with protein misfolding diseases [387].

What is not always clear is to whether these heterotypic interactions tend to promote or to inhibit amyloidogenesis (or even both at different stages, as can occur with the curli protein CsgA and fibrinogen [388]; the same is true for fibrinogen and phenol-soluble modulins [389]).

SERPINA1 (α1 antitrypsin), as found by Schofield et al [202] as a major constituent of microclots, contains four β-sheets and is also able to interact with the amyloidogenic transthyretin [390–392].

Some amyloidogenic proteins, such as Apolipoprotein B-100 that contains a massive crossed-β (‘β-belt’) structure [393], were not detected, however, plausiblty because they were fully embedded within lipoproteins and thus not in plasma. (They are, hopwever, capable of becoming embedded in neurofibrillary tangles [394].) It is therefore approriate next to study

### Amyloidogenicity of proteins ‘entrapped’ in microclots

If axial or lateral co-aggregation is responsible for the ‘entrapment’ of proteins in fibrinaloid microclots, one would suppose that all the proteins involved would themselves be amyloidogenic. This can be tested using amyloidogenicity prediction programs of the type given in Table X. We chose AmyloGram [74], vailable at http://biongram.biotech.uni.wroc.pl/AmyloGram/. For the proteins entrapped within microclots, we took the proteomics data from Table 2 from the Long COVID study of Kruger and colleagues (who did not report on fibrinogen) and the Table 3 of the study of microclots in intensive care patients of Toh and colleagues [202]. The conclusion is very clear: <u>every</u> <u>single one</u> of the proteins detected in the microclots is highly amyloidogenic, and the microclots evidently involve cross-seeding. However, there was little correlation between amyloidogenicity and protein length (Figure 10)(r^2^ = 0.275). However, all but three of the Kruger proteins and all but four of the Schofield ‘top 20’ had an amyloidogenicity score excedding 0.8 (Figure 10).

**Figure 10:**
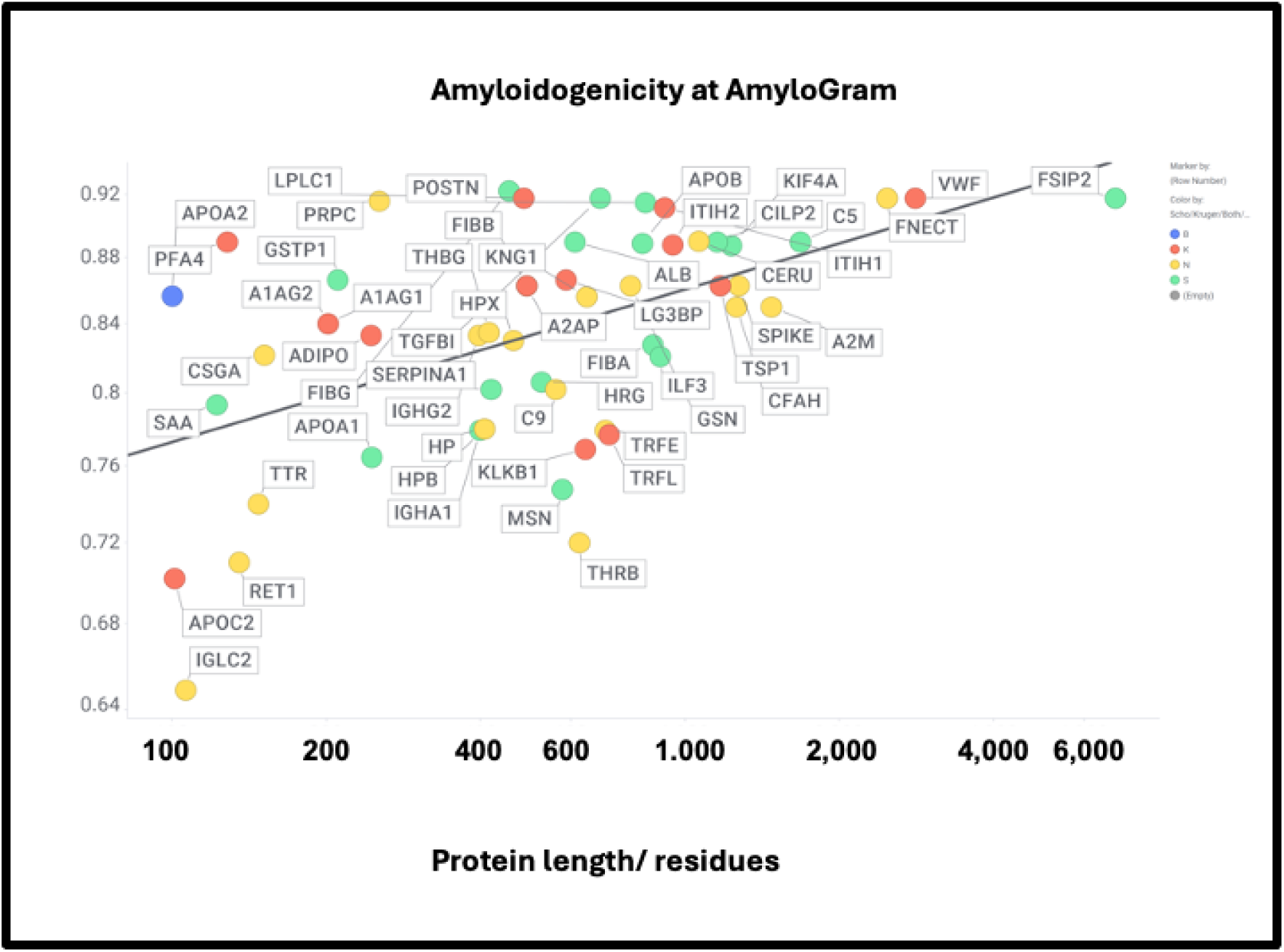
High amyloidogenicity of proteins in the Kruger (red) and Schofield ‘top 20’ (green) studies and in both (blue) plus amyloidogenicity of proteins seen in neither (yellow), and its independence from protein length. The line of best fit indicated has an r^2^ of just 0.23.

**Table 2:** Twelve proteins whose levels differ greatly between fibrinaloid microclots and normal blood clots as seen in proteomics studies (data from. **Figures 9, 10, 11, 16).** Those higher are coded as being from the Kruger (K) [367] or Schofield (S) [202] studies.

| Protein (which study) | Higher or lower in fibrinaloid microclots wrt both normal clots and normal plasma |
| --- | --- |
| Adiponectin (K) | Higher |
| α-2-macroglobulin | Lower |
| Complement Factor 3 | Lower |
| Extracellular Matrix Protein 1 | Lower |
| Factor XIII | Lower |
| Fibronectin | Lower |
| Kallikrein (K) | Higher |
| LBLC1/BNIB1/BNIFB1/LPLUNC1 (K) | Higher |
| Platelet Factor 4 | Higher |
| Periostin (S) | Higher |
| Thrombospondin-1 (K) | Higher |
| Von Willebrand Factor (K) | Higher |

However, there was no correlation with the overall amyloidogenicity as calculated using AmyloGram and either the plasma proteome abundance or whether the proteins were in the fibrinaloid microclots (Figure 11) (r^2^ = 0.02):

**Figure 11:**
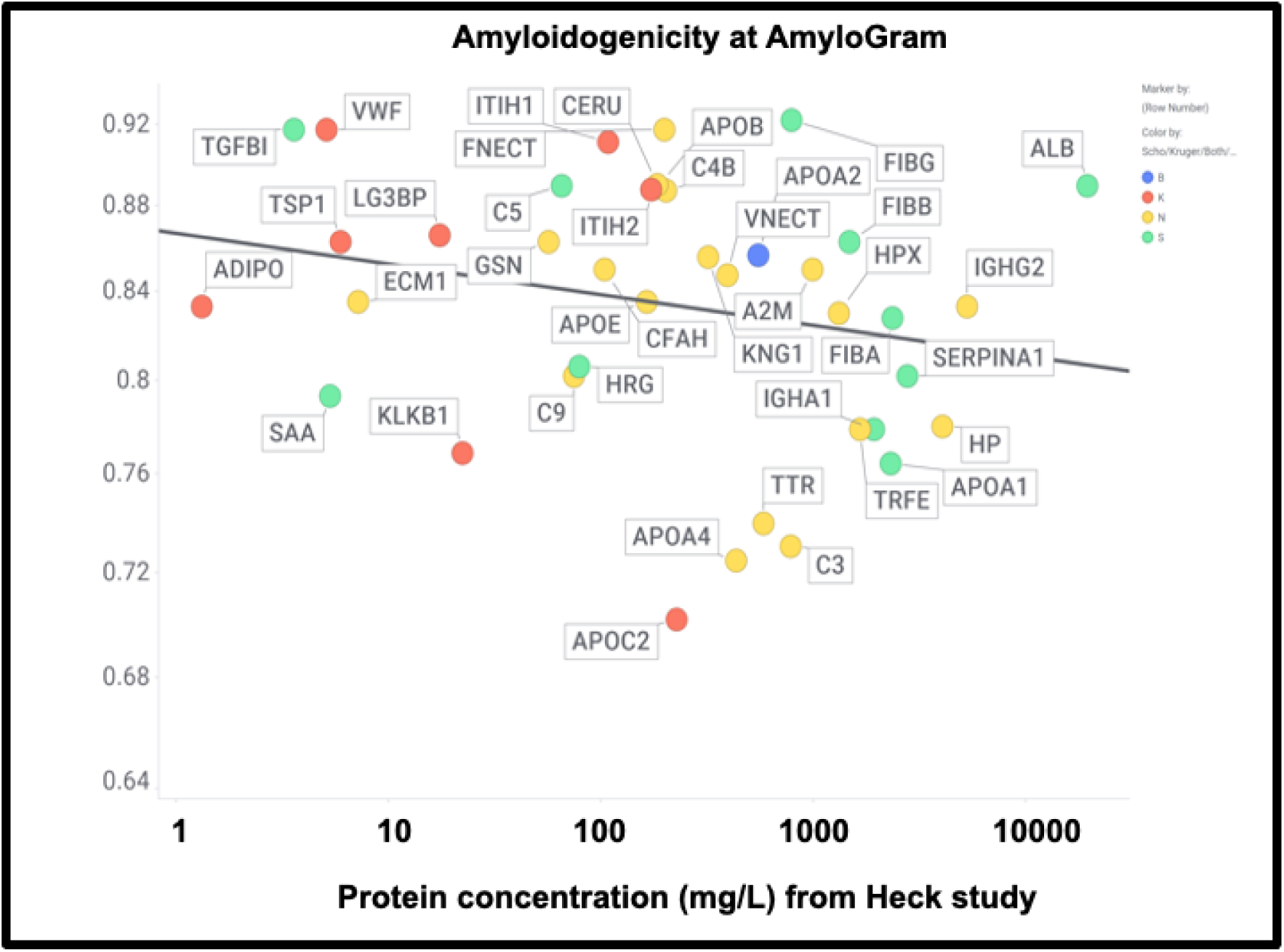
High amyloidogenicity of proteins in the Kruger (red) and Schofield (green) studies and in both (blue) plus amyloidogenicity of proteins seen in neither (yellow), and its independence from plasma protein concentrations slope =0.01, (r^2^=0.05) as recorded in the Heck study [381] (means of first three time points averaged over two controls).

The presence of von Willebrand factor and adiponectin in the fibrinaloid microclots is very inetresting, despite their comparatively low plasma concentration (Figure 11). The former is among the most amyloidogenic proteins in the list (Figure 11), and is notably entrapped and removed by microclots in SARS-CoV-2 infection [395], while the latter is correlated with amyloid Aβ deposition [396] and may be protective [397]. LPLC1 (Fig 10, so low in concentration it does not appear in Fig 11; the human protein atlas https://www.proteinatlas.org/ENSG00000125999-BPIFB1/blood+protein estimates its plasma concentration by mass spectrometry to be 2.7 μg/L) is also of interest. LPLC1 stands for “Long palate, lung and nasal epithelium carcinoma-associated protein 1” or also “BPI fold-containing family B member 1” (BNIB1), and to be clear it is Uniprot Q8TDL5; again it has a very high amyloidogenicity (∼0.91) Finally, thrombospondin-1 is also much over-represented, and it too was neuroprotective against Aβ [383, 398, 399]. A clear pattern emerges.

The precise nature and extent of the amyloidogenicity necessary to induce or be entrapped in fibrinaloid microclots is as yet unclear, but inspection of the detailed data from the analyses at AmyloGram (unpublished) showed that each of the proteins involved possessed a segment of amyloidogenicity (referred to on its website and in the subset of figures displayed here as a ‘probability of self-assembly’) that exceed 0.75 in the data that could be acquired at the AmyloGram website. Figures 12-15 show four examples: the first (Figure 12) is α-2-antiplasmin (prominent in the findings of [303]), where there is an initial run plus two further prominent peaks. α-2-antiplasmin is of course well known as an inhibitor of fibrinolysis [400].

**Figure 12:**
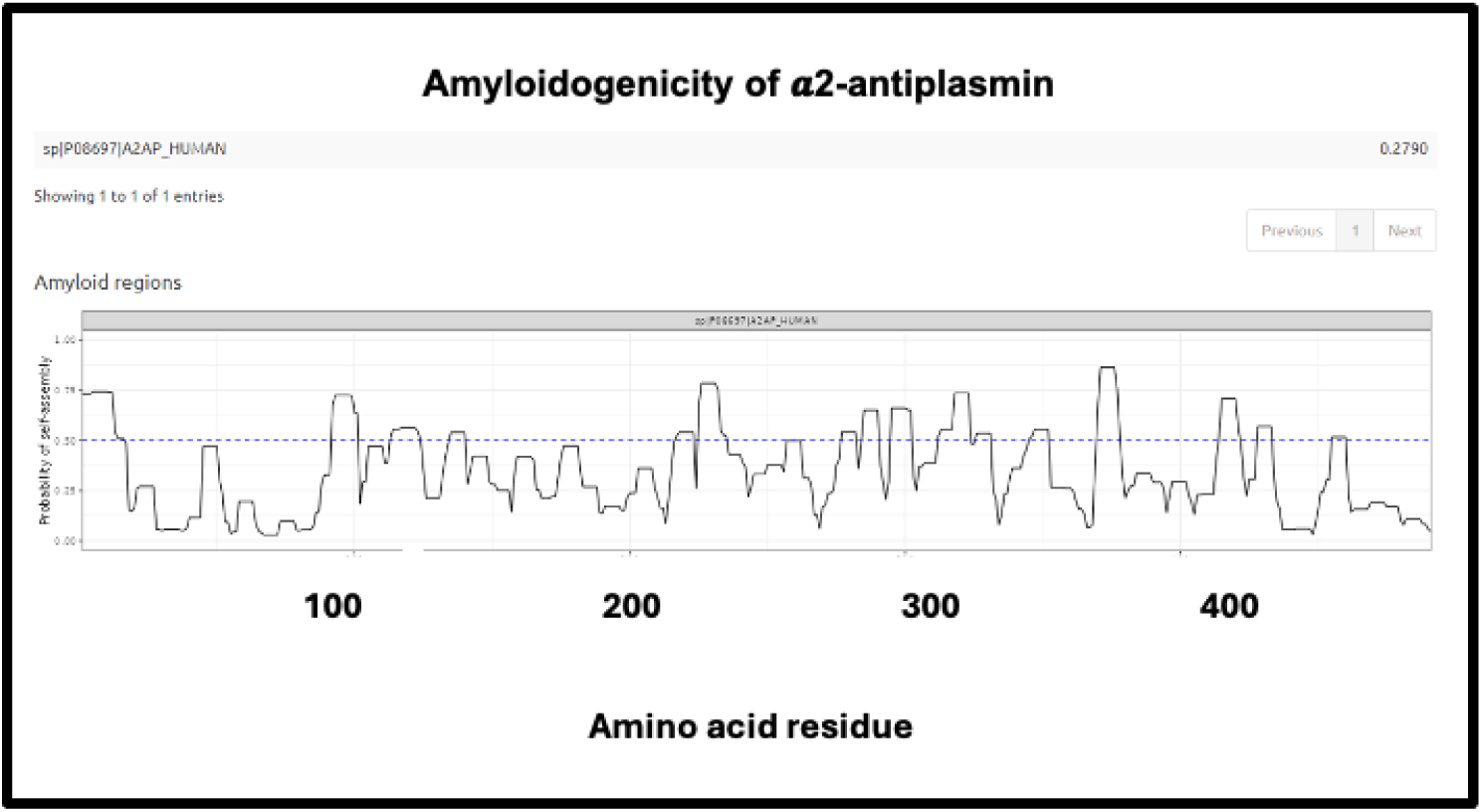
Amyloidogenicity of α-2-antiplasmin.

The next (Figure 13) is SERPINA1 (α1-antitrypsin) where there is a long run at the beginning just exceeding 0.75. SERPINA1 is the most abundant anti-protease in plasma (see also Fig 11), and has several parallel-antiparallel β-sheets in its ground-state conformation [401–405], which, interestingly, is metastable [406–408]. It can also interact with amyloidogenic transthyretin [391], and is associated with the severity and progression of SARS-CoV-2 [409], consistent with its role in assisting fibrinaloid microclot formation. It is thus entirely plausible that it could participate in amyloid formation, and fragments of it certainly do [410, 411].

**Figure 13:**
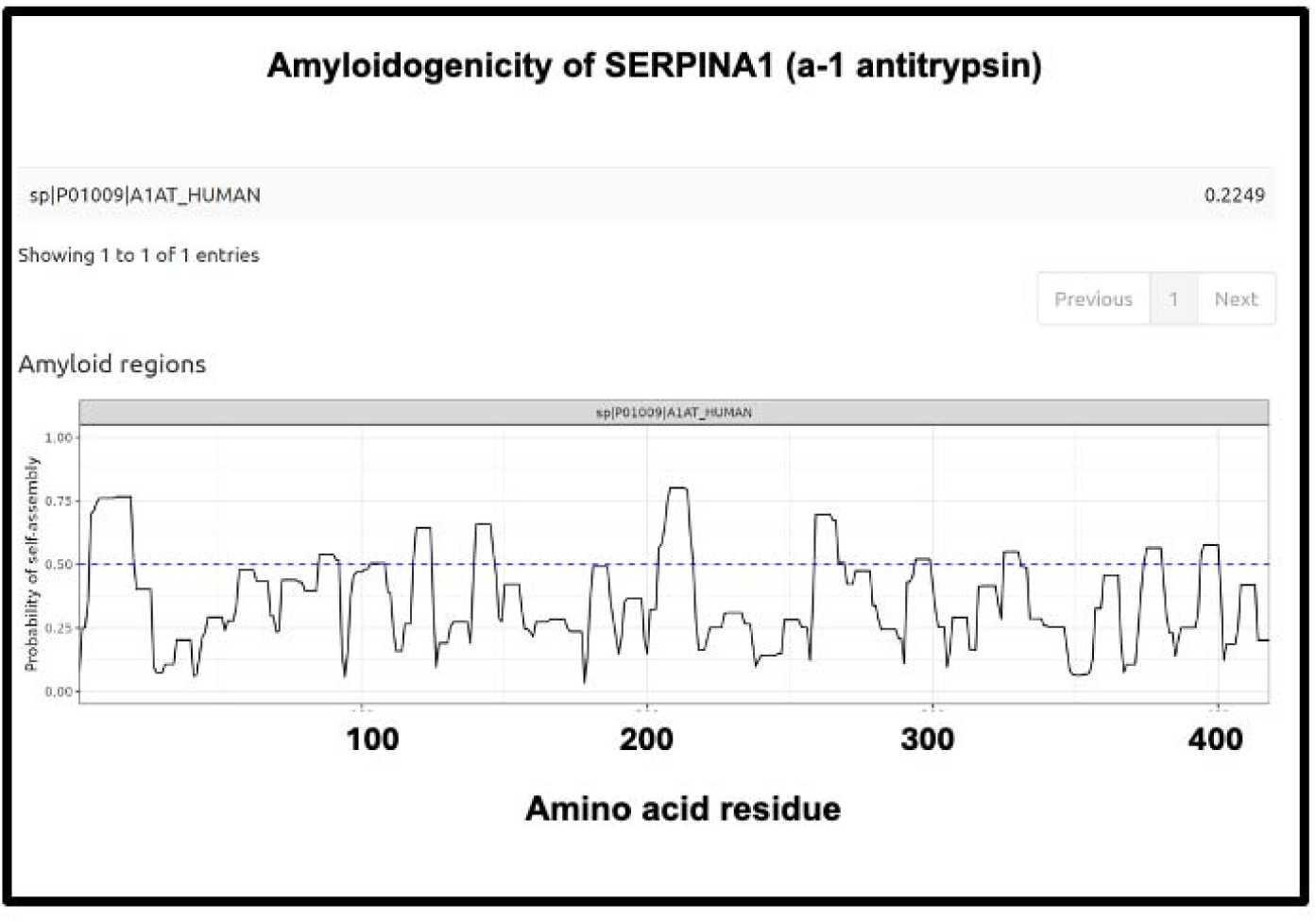
Amyloidogenicity of SERPINA1 (α1-antitrypsin).

**Figure 14:**
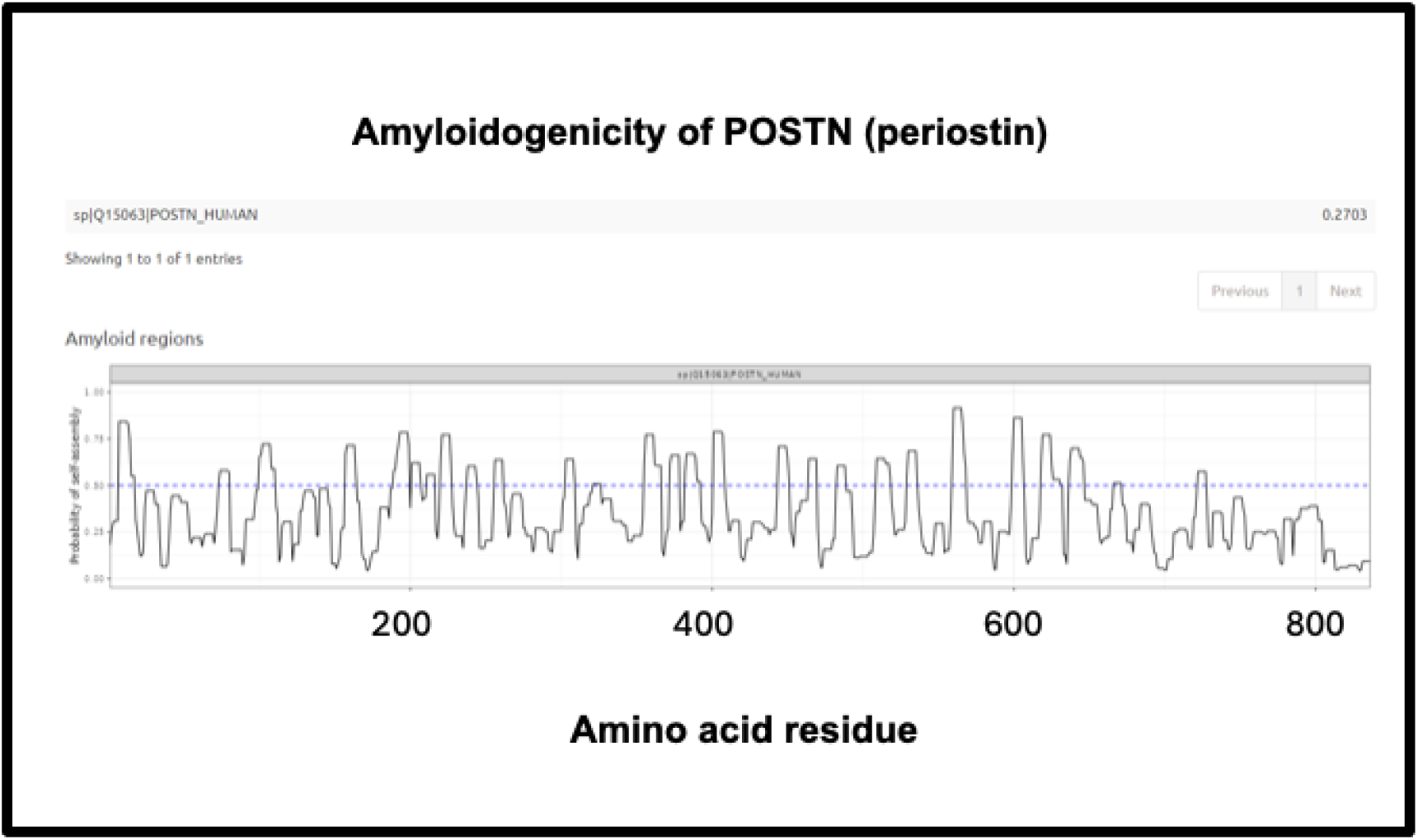
Amyloidogenicity of POSTN (Periostin).

The third is the inflammatory marker periostin, representing 1.17% of the ‘top 20’ proteins in Schofield et al. [202], and containing no fewer than eight regions of amyloidogenicity with a score exceeding 0.75. It features in these fibrinaloid microclots despite being one of the least abundant plasma proteins of those under consideration (372 ng/mL according to [371], 98 ng/mL in [412] and just 10 ng/mL according to [413, 414]) (209 ng/mL is stated for serum [415]). Interestingly, however, as well as being among the most amyloidogenic of those surveyed (Figure 10), it is highly predictive of Aβ deposition [382, 416] and is also involved in lung fibrosis [417, 418].

Finally, the fourth is LPLC1 (also known as BPIB1 or BPIFB1). This protein has a very high amyloidogenic propensity of 0.9176, Figure 10) and no fewer than eight regions in which the amyloidogenicity score exceeds 0.75 (Figure 15), despite a minuscule concentration in normal plasma. Interestingly, it is involved in innate immunity [419], especially in mucosa [420], and bears similarities to lipopolysaccharide-binding protein (LPS being a molecule that can trigger fibrinaloid formation [191, 192, 421]). BPIB1 can also inhibit Epstein-Barr virus proliferation [422–424] (something of major potential relevance in Long COVID [425–427]). Overall, the fact that it is so concentrated in fibrinaloid microclots in the case of Long COVID is thus very notable.

**Figure 15:**
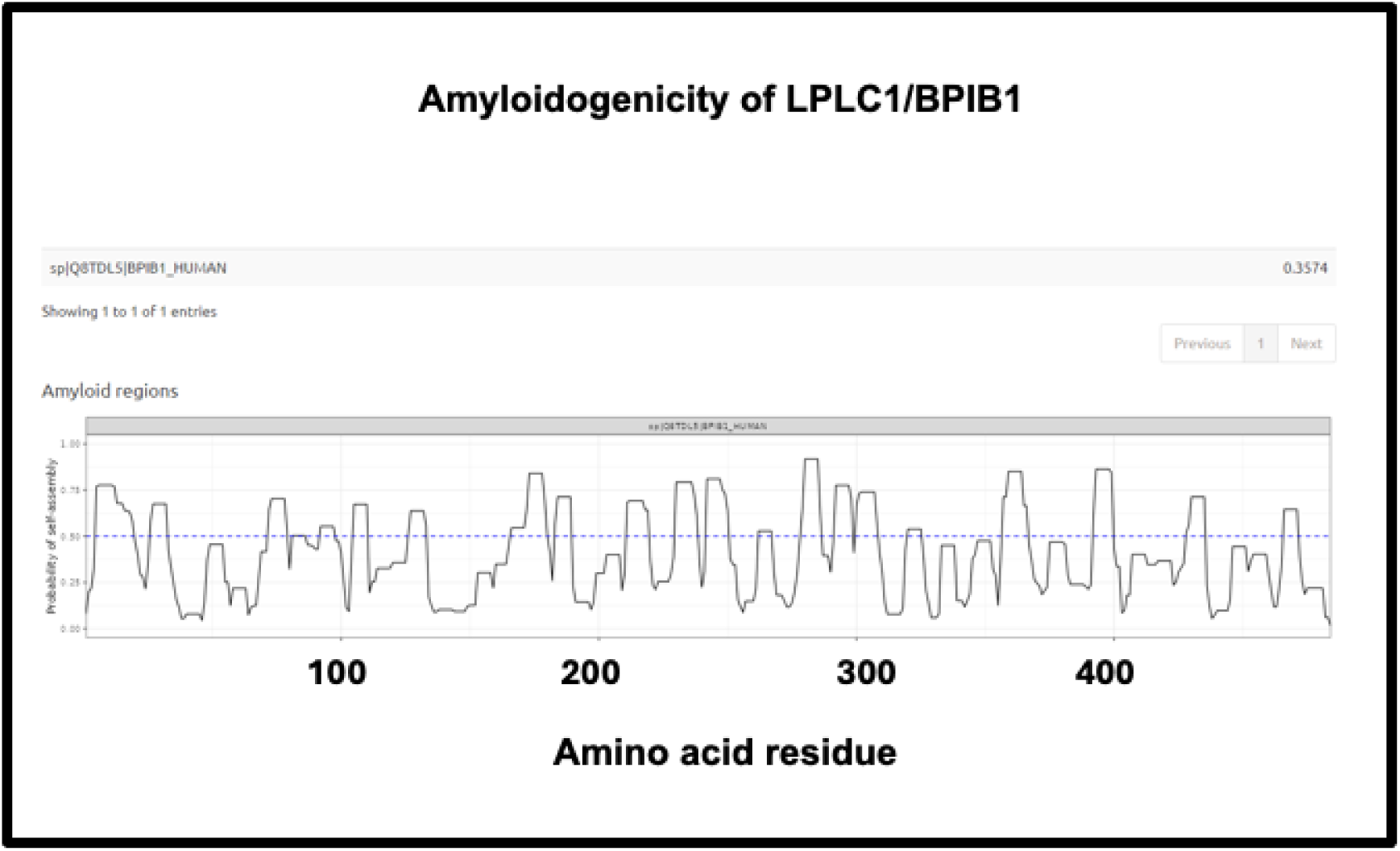
Amyloidogenicity of of LPLC1/BPIB1.

### Comparison with the normal clot proteome

Undas and colleagues [428] provided (their Table 2) a quantitative list of protein-bound proteins in normal clots (that were presumably non-amyloid, though that was not in fact tested). Figure 16 compares the prevalence of proteins in their clots with that of plasma proteins, showing a reasonable correlation (slope = 0.67, r^2^ = 0.29) between the two. This contrasts with that for the fibrinaloid microclot proteome (Figure 7), where the value for r^2^ was just 0.1.

**Figure 16:**
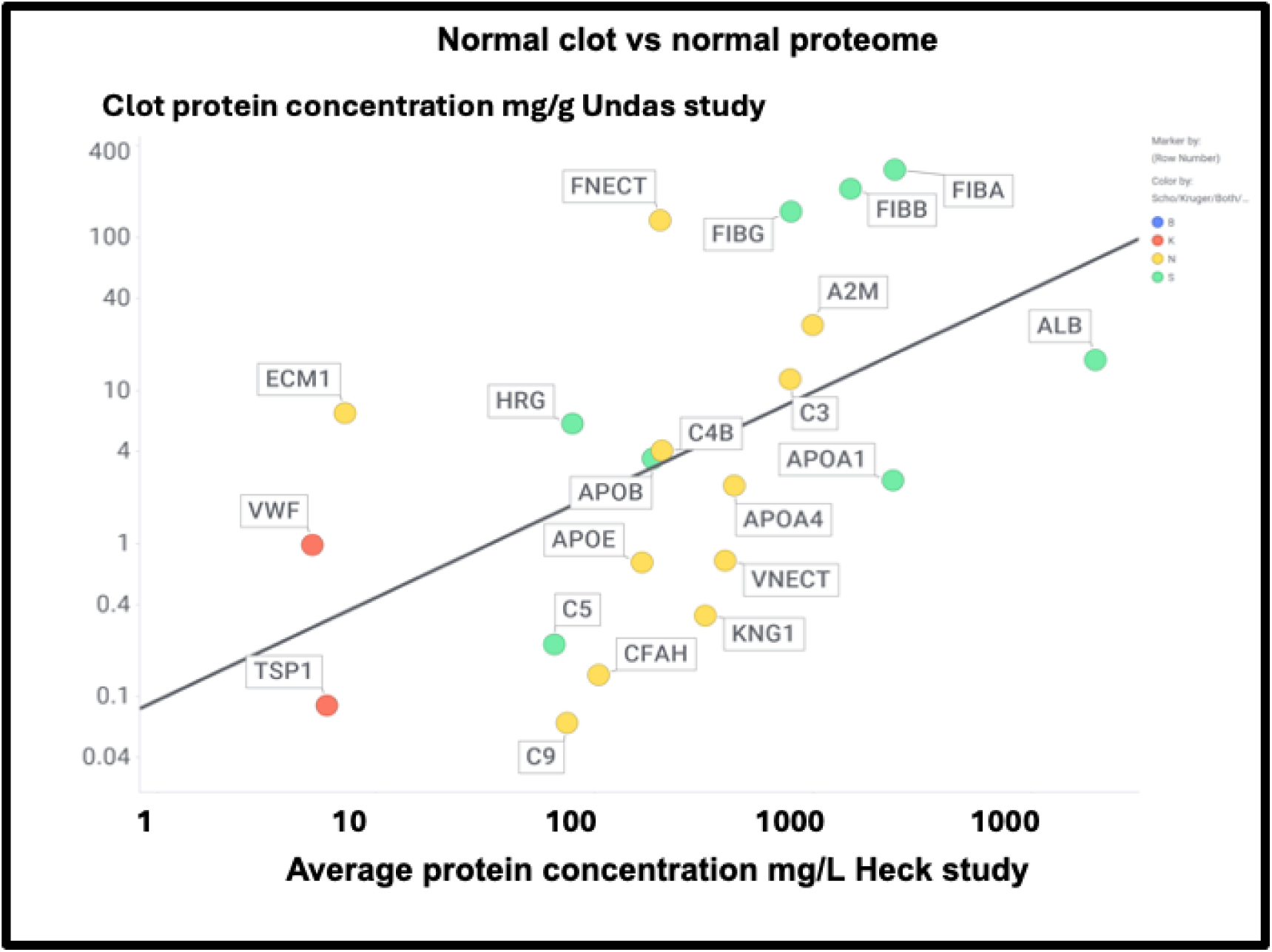
Comparison of the proteome content of normal (non-amyloid) clots vs the standard plasma proteome in controls in the Heck study (average of two controls over first three time points). Colour encoding is as in Figure 11.

Figure 17 shows those that were also assessed in the Kruger and Schofield studies, along with their amyloidogenicity. There is no correlation whatsoever (r^2^ ∼ 0.02), again showing how very different the composition of fibrinaloid microclots is from that of normal clots.

**Figure 17:**
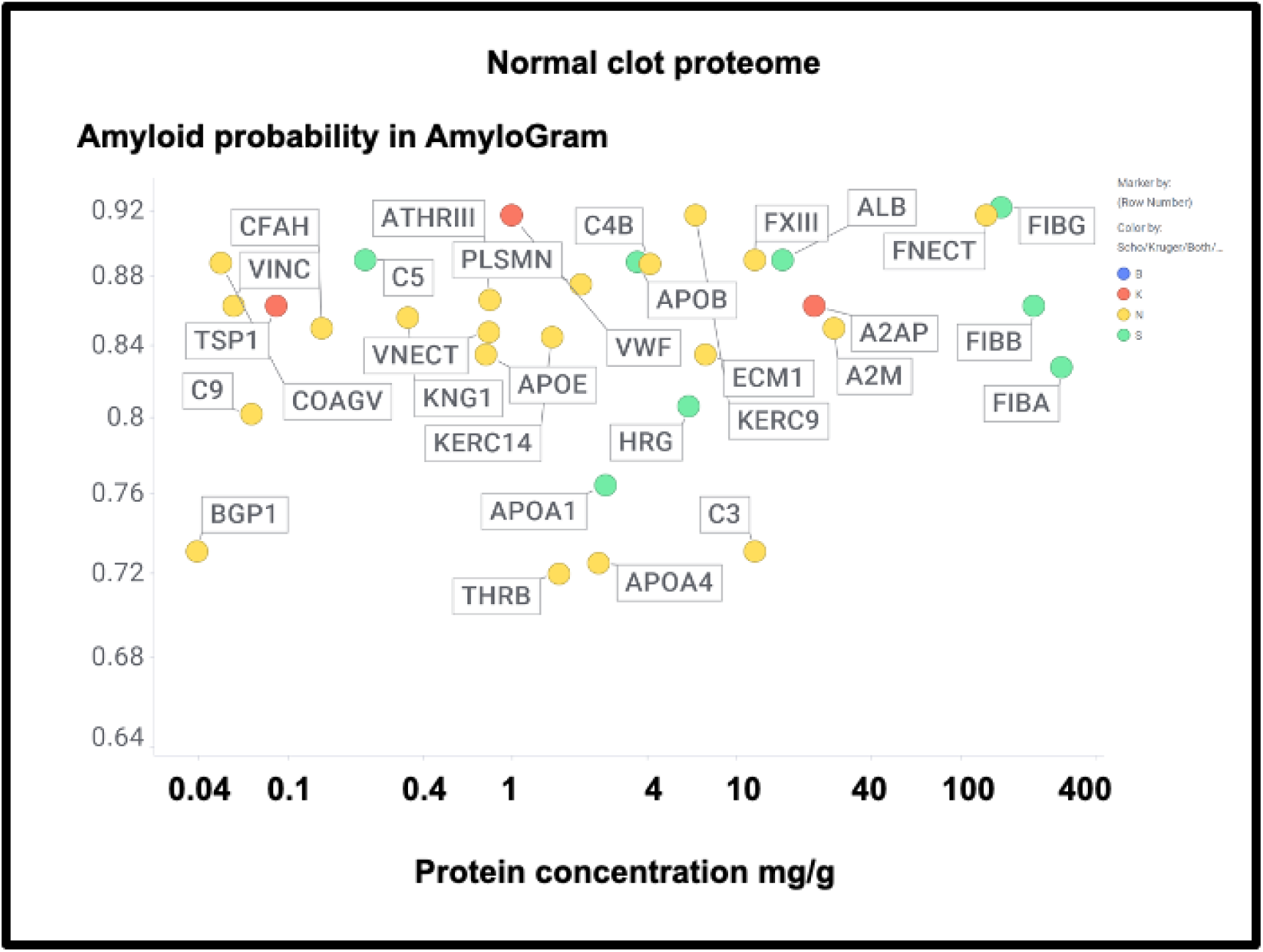
The proteome in normal clots. Data are taken from [428] and also coded as to whether the proteins were observed in the fibrinaloid microclots observed by Schofield (green), Kruger (red), both (blue) or neither (yellow).

A particularly noteworthy observation here (Figure 17) is the relatively high amounts of fibronectin seen in normal clots (130 mg/g protein [428], fibronectin typically being present in plasma at 300-400 mg/L [429–431]) as it was <u>not</u> seen in the fibrinaloid microclots. While fibrin is highly amyloidogenic (Figures 10, 17) and *in vivo* can produce insoluble fibrillar components that may be incorporated in the extracellular matrix [432–434], fibronectin is somewhat unusual for two reason. First, it is large (2477 residues). Secondly it is relatively thermostable [435], especially in some of its domains [436]. Together these features can plausibly account for the difficulty of unfolding and incorporating it into an amyloid clot compared to a normal one (and see also next paragraph for amyloidogenicity vs thermostability). Similar comments relate to α-2-macroglobulin (27 mg/g, 1474 residues, Figure 17) and to Factor XIII (12 mg/g, 732 residues), which is in fact inhibited by α-2-macroglobulin. Factor XIII is a transglutaminase (linking glutamate and lysine residues) that is responsible for stiffening normal clots by crosslinking them [437–439], mainly via the γ but also partly the α chains [440], yet does not appear in the fibrinaloid microclot proteomes. This would be entirely consistent with their completely different structures relative to that of normal clots. Lastly, complement factor 3 is fairly well represented in both plasma (Figure 16; 778 mg/L in the Heck study) and in normal clots (Figure 17; 12 mg/g) yet is not found in fibrinaloid microclots; consistent with the thrust of our arguments, its total amyloidogenic propensity at just 0.73 is among the lowest of those evinced in this study.

### Amyloidogenicity vs thermostability

As noted, amyloidogenesis of a non-amyloid form of a protein necessarily requires a significant unfolding, and in general terms the resistance to a protein unfolding is reflected in its thermostability.The point about the experimental lack of amyloidogenicity in a thermostable protein thus leads to the question of of whether experimental amyloidogenic proteins are in fact normally relatively non-thermostable. This turns out to be strongly supported by substantial evidence [249, 258, 441–447].

## Discussion

A general question about amyloid(ogenic) protein fibres in general, and fibrinaloid microclots (our main focus) in particular, is the nature and location of the proteins that they contain. A variety of studies have provided data on both normal clots and fibrinaloid microclots, as well as the normal plasma proteome. With occasional exceptions (such as C-reactive protein – not involved here) their concentrations are fairly constant, and since (i) they cover four orders of magnitude in the proteins considered here, and (ii) they were well correlated in two studies (Figure 8), we consider their standard concentrations to be a good guide as to the likelihood of non-fibrinogen proteins being entrapped in a clot if entrapment simply reflects their plasma concentrations. In normal clots that expectation is broadly borne out (Figure 15). However, this is <u>far</u> from being the case with the fibrinaloid microclots that form in certain diseases, stain strongly with thioflavin T, and are far more resistant than are normal clots to fibrinolysis (Figure 18). Seemingly, as with prion proteins, the presence of a small amount of the thermodynamically stabler amyloid form is enough to trigger conversion of a very large number of monomers in the amyloid polymer form in almost an ‘all-or-nothing’ manner (Figure 19)

**Figure 18:**
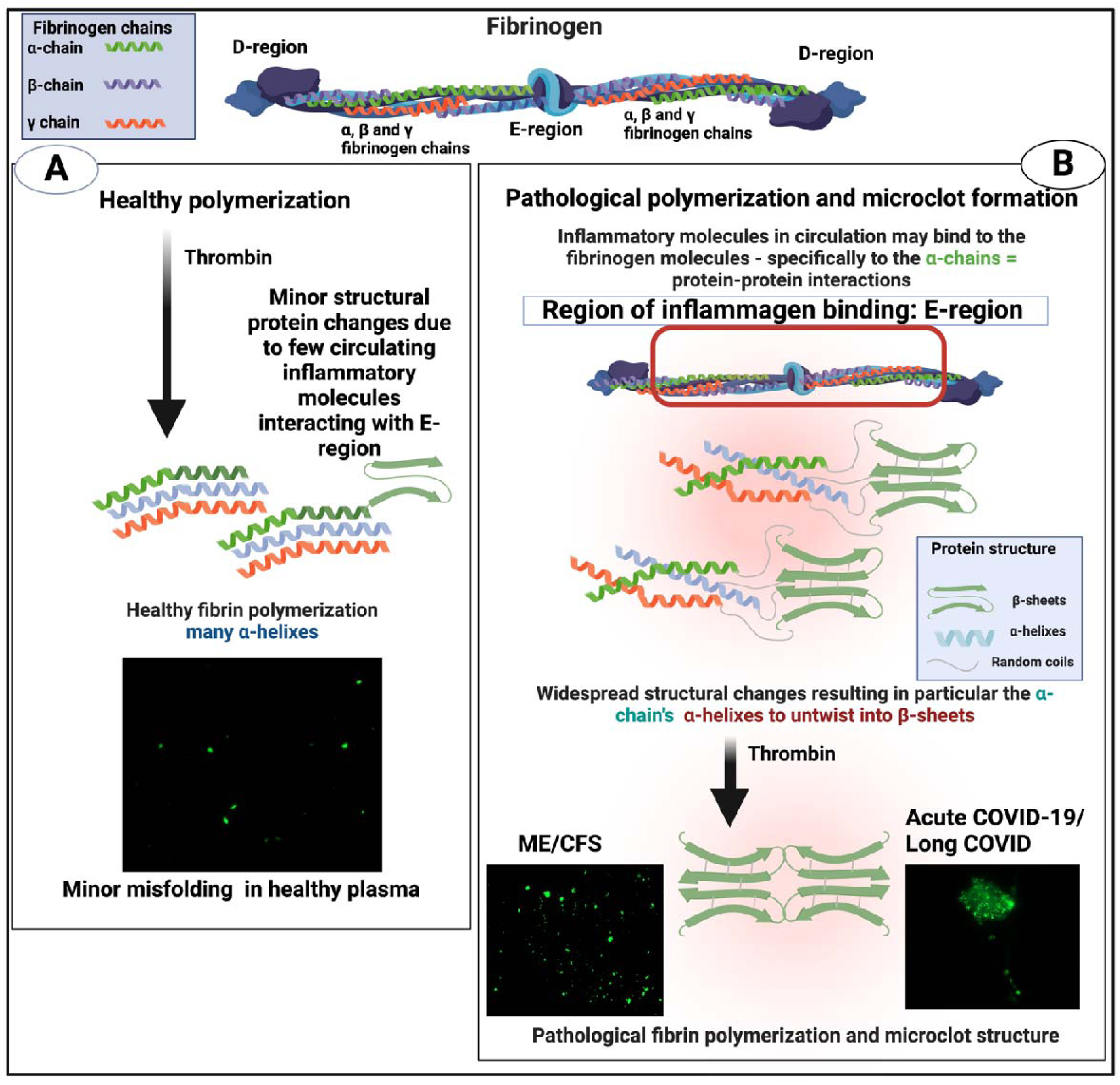
Healthy versus pathological (amyloid) clotting Taken from [22]). Created with Biorender.com.

**Figure 19:**
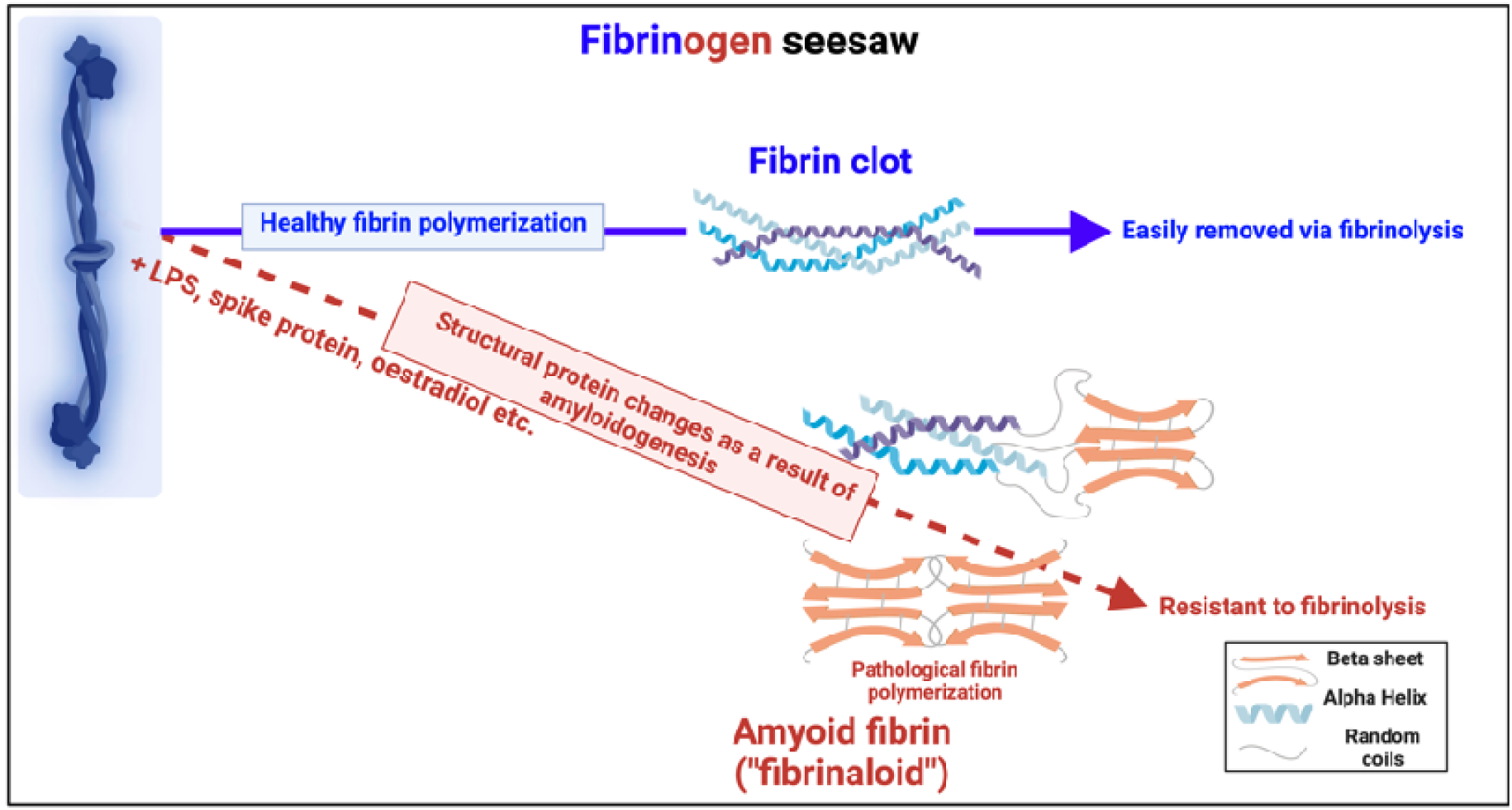
Individual fibrinogen molecules upon polymerisation either polymerise into a normal clot form, that is relatively easily removed by fibrinolysis, or into an anomalous amyloid form or forms, that are not (Taken from [449]). Created with Biorender.com.

First, their proteome varies strongly with the disease (Figure 7), in a way that cannot reflect changes in the proteome. Secondly, their proteome constitution is far from being related to the concentrations of bulk plasma proteins, with some being excluded and others being highly concentrated (Figures 9, 10, 11, 16; twelve are summarised for convenience in Table 2). Clearly there must be special mechanisms at work, the most obvious, given their strikingly high amyloidogenicity scores, being a cross-seeding where the various proteins are actually incorporated into the cross-β elements of the fibrils themselves. This said, there is a consonance in Table 2 between the proteins highlighted as being in fibrinaloid microclots and biological explanations based on their known roles. Most of those that are higher are in the Kruger [367] Long COVID study (Table 2). Long COVID is of course a chronic disease, and very different from the acute conditions characterising individual in an ICU such as those in the Toh study [202]. It bears relations to myalgic encephalopathy/chronic fatigue syndrome (ME/CFS), however, so it is of interest that thrombospondin and platelet factor 4 are both raised in the plasma of individuals with ME/CFS [448].

### Clot fibrinaloids vs classical amyloid fibrils

Much is known about classical amyloids, and plausibly this knowledge will contribute to our emerging understanding of fibrinaloids, but there is one crucial difference and that is the size of the fibres. A typical oligomer made up of a standard cross-β amyloid chain crisscrossing in a direction perpendicular to the fibril direction has a diameter in the range 2 to 5 nm [450], and these can aggregate to make protofilbrils in the range 4-11nm diameter [451]. The amyloid fibrils themselves typically involve a few intertwined protofibrils per unit length [270], and thus have a diameter of the order of 6 to 12 nm [452–454], although in principle they could become larger [270]. This means that generally most fibrils consists of only a few intertwined protofilbrils per unit length. By contrast, the fibrils observed in fibrinaloid microclots in plasma samples have a diameter that is often in the range 50 to 100 nm. This means that they must contain many more lateral fibres per unit length, likely of the order of 100-300. This ability to increase numbers laterally obviously bears strongly on the potential ability of microclots to aggregate further to become macroclots, and maybe ultimately to lead to the kinds of occusions involved in stroke and myocardial infarctions.

Another particular feature of the fibrinaloid microclots is their resistance to fibrinolysis (see [21, 22, 195, 303, 455, 456], most easily seen in our proteome studies where a <u>double</u> trypsin digestion was required for successful peptide-based mass spectrometric proteome analysis [303, 367]. The presence of molecules such as α-2-antiplasmin [303] and SERPINA1 will certainly have contributed, but it is of course well known that amyloid form of proteins are far more resistant to proteolysis than their native unfolded or globular versions, especially among prion proteins where PrP^Sc^ is even resistant to proteolysis by proteinase K (e.g. [457–461]).

Some molecules such as bacterial cell wall compounds can clearly serve as triggers for amyloidogenesis in both fibrinaloid microclots [191–193, 421, 462–465] and other cases [466–468]. What we propose here is that the massive changes in fibrinogen structure necessary for its conversion to an amyloid form, as also oberved by others [201, 202, 469], must then involve cross-seeding, as this provides a simple mechanism that at once accounts for (i) the proteomics, (ii) the resistance to proteolysis, and (iii) the amyloid nature of the mixed clots. At the same time, we recognise that while our analysis is both clear and robust, future studies would benefit from comparing the proteomes of fibrinaloid and normal clots at the same time on the same instrument.

Overall, however, while the present findings make it clear that cross-seeded proteins must be involved in firbrinaloid formation, we are far from knowing their specific locations and whether and how they co-aggregate axially and/or laterally (as per Figure 6). Such studies would require the resolution of the atomic force microscope (e.g. [223, 470]), which on the basis of the present findings it now seems worthwhile to pursue.

## Materials and methods

Data were downloaded from the sources indicated, and were not otherwise transformed save to average the first three timepoints and two individuals representing the controls in [381]. The values from Figure 3 of Schofield *et al*. [202] were determined digitally from the averaged pie chart therein. An Excel file provided as Supplementary Information summarises the data, that are commonly displayed in this paper using the Spotfire program (https://www.spotfire.com/).

## Author Contributions

Both authors contrubted to the conceptualisation, analyses, funding acquisition, drafting, and final editing.

## Acknowledgments

DBK thanks Prof Ben Goult for useful discussions.

## Abbreviations

A1AG1: alpha-1-acid glycoprotein-1
A1AG2: Alpha-1-acid glycoprotein 2
A2AP: alpha-2 antiplasmin
A2M: Alpha-2-macroglobulin
ADIPO: adiponectin
ALB: Albumin
APOA1: Apolipoprotein A-1
APOA2: Apolipoprotein A-II
APOA4: Apolipoprotein A-IV
APOB: Apolipoprotein B
APOC2: apolipoprotein C-II
APOE: Apolipoprotein E
ATHRIII: Antithrombin-III
BGP1: Beta-2-glycoprotein 1
C3: Complement C3
C4B: Complement C-4B
C5: complement C5
C9: Complement C9
CERU: Ceruloplasmin
CFAH: Complement Factor H
CILP2: cartilage intermediate layer protein 2
COAGV: Coagulation Factor V
CSGA: Curli protein E coli
ECM1: Extracellular matrix protein 1
FIBA: fibrinogen alpha chain
FIBB: Fibrinogen beta chain
FIBG: Fibrinogen gamma chain
FNECT: Fibronectin
FSIP2: Fibrous sheath-interacting protein 2
FXIII: Factor XIII A chain
GSN: Gelsolin
GSTP1: glutathione S-transferase P
HP: Haptoglobin
HPB: Haptoglobulin beta chain
HPX: Hemopexin
HRG: histidine-rich glycoprotein
IGHA1: immunoglobulin heavy constant alpha 1
IGHG2: Ig gamma 2 heavy chain
IGLC2: Ig lambda light chain
ILF3: interleukin enhancer-binding factor 3
ITIH1: Inter-alpha-trypsin inhibitor heavy chain H1
ITIH2: Inter-alpha-trypsin inhibitor heavy chain H2
KERC14: Keratin type 1 cytoskeletal 14
KERC9: Keratin type 1 cytoskeletal 9
KIF4A: KIF4A
KLKB1: kallikrein
KNG1: kininogen
LG3BP: Galectin-3-binding protein
LPLC1: Long palate, lung and nasal epithelium carcinoma-associated protein 1
ME/CFS: myalgic encephalopathy/chronic fatigue syndrome
MSN: Moesin
PFA4: Platelet basic protein/Platelet factor 4
PLSMN: Plasmin(ogen)
POSTN: Periostin
PRPC: Human prion protein
RET1: Retinol-binding protein
SAA: serum amyloid A
SERPINA1: SERPINA1
SPIKE: SARS-CoV-2 spike
TGFBI: transforming growth factor-beta–induced protein ig-h3.
THBG: Thyroxine binding globulin
THRB: (Pro)thrombin
TRFE: transferrin
TRFL: lactotransferrin
TSP1: thrombospondin-1
TTR: Transthyretin subunit
VINC: Vinculin
VNECT: Vitronectin
VWF: von Willebrand Factor

